# Efficient value encoding through convergence of tactile and visual value information in the primate putamen

**DOI:** 10.1101/2024.01.04.574258

**Authors:** Seong-Hwan Hwang, Doyoung Park, Ji-Woo Lee, Sue-Hyun Lee, Hyoung F. Kim

## Abstract

The processing of diverse sensory values by a limited number of basal ganglia neurons raises the question of whether each value is processed independently or combined as the number of neurons decreases from the cortex to downstream structures. Here, we discovered that tactile and visual values were partially converged in the primate putamen, enhancing its efficiency in encoding values while preserving modality information. Humans and monkeys performed tactile and visual value discrimination tasks. Notably, the human putamen selectively represented both tactile and visual values in fMRI scans. Single-unit electrophysiology further revealed that half of the individual neurons in the macaque putamen encoded both tactile and visual values, and the other half encoded each value separately. The bimodal value neurons enable more efficient value encoding using fewer neurons than the modality-selective value neurons. Our data suggest that the basal ganglia system uses modality convergence to efficiently encode values with limited resources.

## Introduction

Various types of values can originate from sensory input such as tactile and visual sensations. The basal ganglia (BG) might process values from these diverse sensory inputs because neurons in the cortical regions that process each type of sensory information innervate structures in the BG system^1–3^. Interestingly, the number of neurons is known to decrease as information flows from the cortex to each successive structure in the BG^4–6^. This anatomical observation raises the question of whether each value from distinct sensory inputs is processed independently to retain both value and modality information or is combined to preserve only value information. This question can be explored through opposing classical hypotheses that persist in an ongoing debate and remain unresolved: parallel information processing and convergent information processing^4–9^.

Primates use their hands and fingers to perceive tactile input and use this sensory information in their decision-making process^10,11^. For example, humans can locate a dropped coin among various objects beneath a sofa using tactile perception through their fingers. Similar searching behavior is commonly observed among primates as they skillfully use their finger perception to sift through and select valuable objects placed inside various cavities or crevices^12–15^.

This foraging behavior with finger exploration provides insights into the cognitive processes through which primates discriminate tactile stimuli, remember their values, and make decisions based on their finger perception. However, little research has investigated the brain regions responsible for processing tactile value information. Most research has focused on cognitive processes related to visual values in the primate brain. To complete tasks involving visual stimuli, structures in the BG system, including the putamen and caudate, process the value information^16–19^. Notably, the putamen in the BG system anatomically receives input from the somatosensory cortex, including finger regions, and from midbrain dopamine neurons for value encoding^3,20,21^. That suggests a plausible role for the putamen in processing value information about tactile stimuli perceived through the fingers. Consequently, it is conceivable that neurons in the putamen are responsible for processing value information not only from the visual modality but also from the tactile modality. Investigating the convergent or parallel processing of tactile and visual value information in the putamen is key to understanding how the BG system deals with the decreasing number of neurons as information progresses from the cortex to successive structures.

In this work, we examined the striatal regions using functional magnetic resonance imaging (fMRI) and discovered that the human putamen selectively represents both tactile and visual value information. To further understand how individual neurons process these different values within a single structure, we conducted single-neuron recordings in macaque brains while the monkeys performed both tactile and visual value discrimination tasks. In this study, our focus was on the striatum, a structure that receives information from cortical areas and is known to encode visual value information^16,22^. We investigated whether the processing of tactile and visual values uses convergent or parallel processing.

## Results

### Representation of both tactile and visual values in the human putamen

To identify which striatal structures are involved in processing tactile and visual values and how they function, we conducted a human fMRI experiment with two value reversal tasks: a tactile task and a visual task (Fig. 1). Participants experienced both tactile (braille) and visual stimuli (fractal image) in each trial of the value reversal tasks (Fig. 1a). In the tactile task, monetary gain or loss was associated exclusively with the identity of the tactile stimulus, and in the visual task, it was associated only with the identity of the visual stimulus (Fig. 1b). After experiencing these stimuli, participants were instructed to press a button to indicate whether the stimulus was associated with a good or bad monetary reward, and then they received feedback about their response (Fig. 1c). To distinguish brain responses for value processing from those for stimulus identity processing, the stimulus–reward contingency was reversed in the middle of each task run (Fig. 1b). We found that participants successfully learned the stimulus–value associations and made responses aimed at maximizing their total monetary gain in both the tactile and visual tasks (tactile task: 96.59 ± 0.59% accuracy; visual task: 96.95 ± 0.64% accuracy; excluding the first trial and the rule-switching trial in each run). There was no significant difference in accuracy between the two tasks (t(21) = −0.458, p = 0.652), indicating the acquisition of a comparable level of value information from both sensory modalities used.

**Figure 1.**
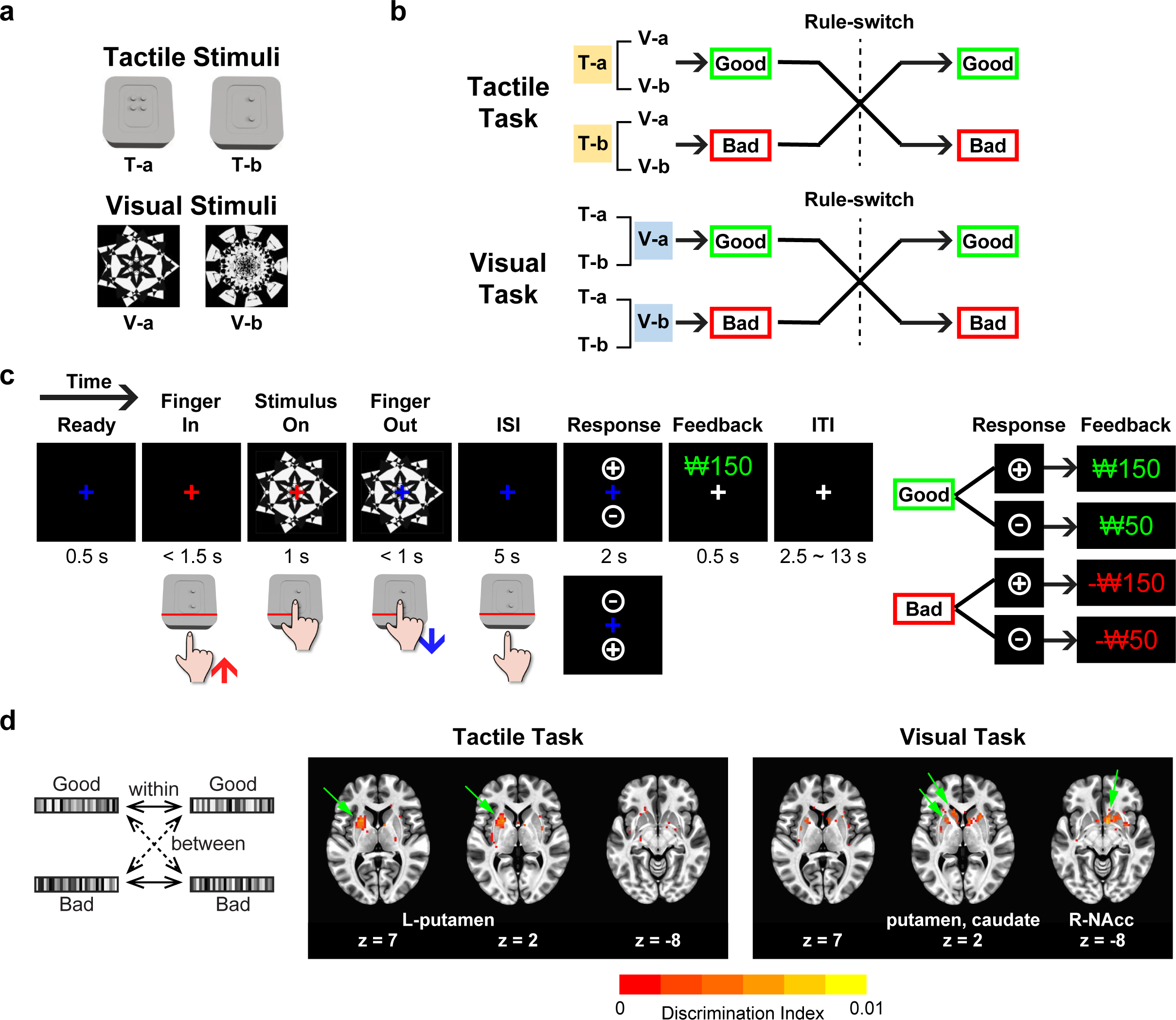
Selective representation of both tactile and visual values in the human putamen. **(a)** Tactile and visual stimuli for human participants. Two braille patterns and two fractal images were used as the tactile and visual stimuli, respectively. **(b)** Value reversal scheme. In each task, either tactile or visual stimuli were associated with monetary values, and stimuli from the other modality were devoid of monetary associations. Within each task run, one stimulus was linked to monetary gain, and the other was paired with monetary loss. This stimulus–reward contingency was reversed in the middle of each run of both tasks. **(c)** Tactile and visual value reversal tasks. Participants were presented with both tactile and visual stimuli simultaneously, with one modality being associated with a monetary gain or loss according to the task. Participants were instructed to press a button indicating whether the stimulus was associated with a good or bad monetary reward. Correct responses to stimuli associated with a monetary gain earned participants 150 won, and incorrect responses provided 50 won (good stimuli). In contrast, correct responses to stimuli associated with a monetary loss deducted 50 won, and incorrect responses deducted 150 won (bad stimuli). **(d)** Searchlight results showing both tactile and visual value representations in the putamen. The discrimination index for value information was calculated as the difference between the averages of within-value correlations and between-value correlations. The colored areas indicate significant discrimination of value across participants (p < 0.05, t_(21)_ > 1.721) delineated on axial slices of the MNI brain.

Next, we examined which striatal structures represented each type of value information based on the human BOLD responses. Using a searchlight procedure, we identified structures with positive value discrimination indices. These indices were computed based on the mean voxel pattern correlation between trials with the same value (e.g., good-and-good or bad-and-bad) subtracted by the mean correlation between trials with opposite values (good-and-bad) (Fig. 1d, left panel)^23,24^. Notably, our searchlight analysis revealed a significant value discrimination response in the putamen of the left hemisphere during the tactile task, corresponding to the side contralateral to tactile perception by the right fingertips (Fig. 1d, middle panel). During the visual task, we found a significant value discrimination response in the bilateral putamen, as well as in the caudate and right ventral striatum (Fig. 1d, right panel). These results demonstrate that the human putamen selectively represented both tactile and visual values, implying the potential for value-convergent processing in the human putamen.

### Learning the values of tactile and visual stimuli in macaque monkeys

The human fMRI study revealed that both tactile and visual values were represented in the putamen, raising the question of whether a single neuron converges both values within this structure. To explore how single neurons in the putamen process tactile and visual value information, we designed a tactile value reversal task (T-VRT) and visual value reversal task (V-VRT) using a newly developed tactile stimulus presenter for macaque monkeys (Figs. 2a–d, and S1). The two monkeys (UL and EV) were required to insert their fingers into the finger hole to experience a braille pattern as a tactile stimulus in the T-VRT (Figs. 2a and S1b-c) (Movie S1). Similarly, a fractal image was presented on the screen when the monkeys inserted their fingers into the hole in the V-VRT. In each block of these tasks, one tactile or visual stimulus was associated with a reward (good stimulus), and the other stimulus was not (bad stimulus) (Fig. 2b). This stimulus–reward association was reversed in each block to investigate neural responses encoding values while excluding neural responses to the stimuli themselves.

**Figure 2.**
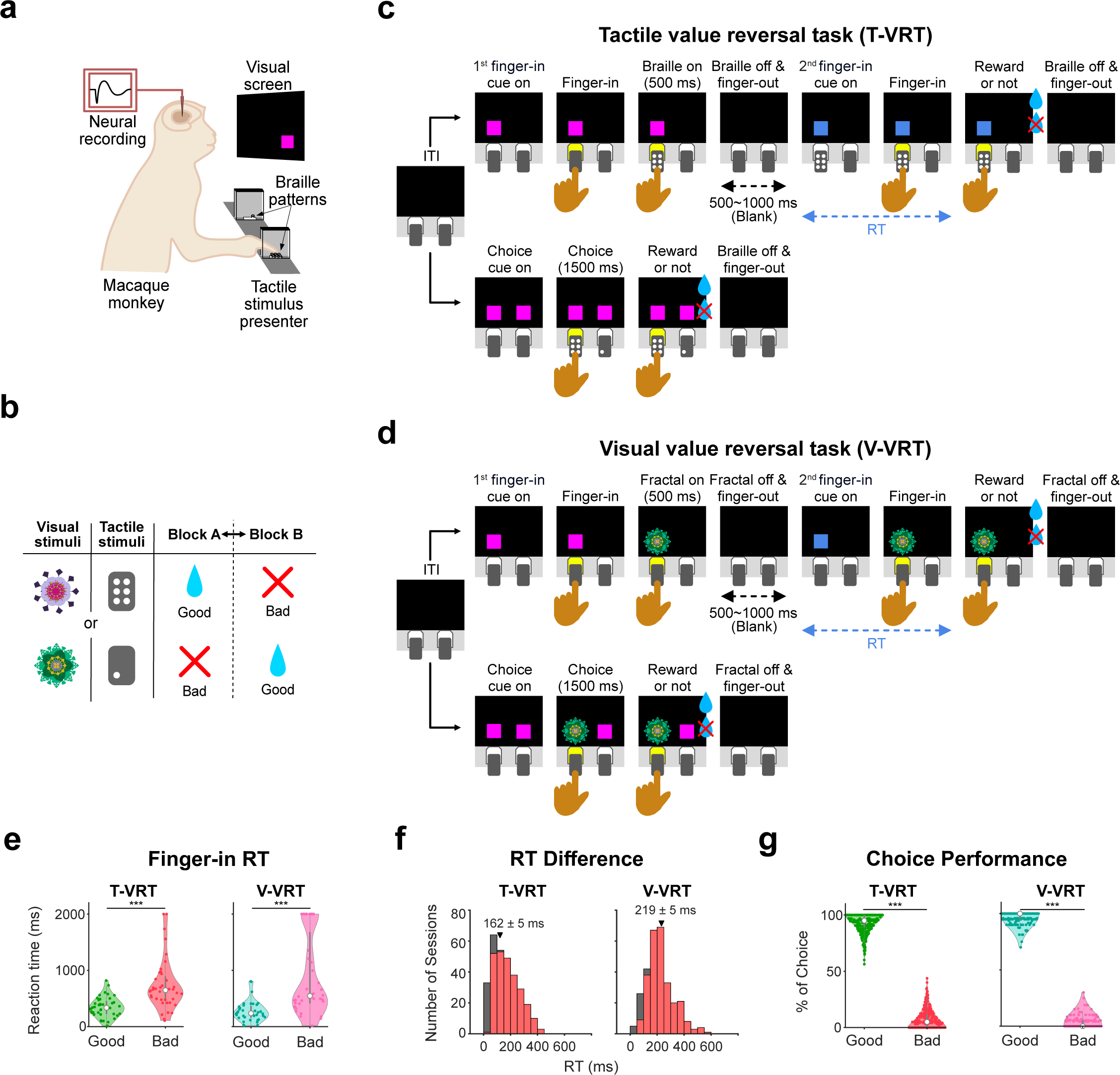
Tactile and visual value learning in macaque monkeys. **(a)** Schematic of the experimental conditions. Tactile and visual stimuli were presented through the braille presenter and monitor screen, respectively. Each monkey’s head was fixed for electrophysiology and viewing visual stimuli on the screen. **(b)** Reversal of value for tactile and visual stimuli. In each block, either a braille or fractal stimulus was associated with a liquid reward, and the other was not. This stimulus-reward contingency was then reversed in the following block. **(c)** Tactile value reversal task (T-VRT). The T-VRT includes two types of trials: single-stimulus trials and choice trials. In the single-stimulus trials, the monkeys experienced a single tactile stimulus and received the associated reward. In the choice trials, two squares for selection were presented. **(d)** Visual value reversal task (V-VRT). The procedure for the V-VRT was identical to that for the T-VRT except for the modality of the stimuli. **(e)** Example of faster reaction times for finger insertion for good stimuli than bad stimuli. The monkeys inserted their fingers more quickly for anticipated good stimuli than bad stimuli in response to the second finger-in cue (n = 40 and n = 36 for the T-VRT and V-VRT, respectively). **(f)** Histogram of averaged differences in finger insertion reaction times measured across all the recording sessions for the two monkeys (n = 299 for both the T-VRT and V-VRT). Red bars indicate neurons with statistically significant differences (two-tailed unpaired t-test, p < 0.05). Triangles and numerical values indicate the mean differences from all sessions (mean ± SE). **(g)** Percentage of choices in the choice trials. The percentages for selecting good and bad stimuli were plotted (n = 299 for both the T-VRT and V-VRT).

To evaluate whether monkeys learned the tactile and visual values in these tasks, we first examined the differences in finger insertion reaction times (RTs) when they anticipated a good or bad stimulus. Each stimulus was presented twice within a single-stimulus presentation trial (Figs. 2c–d, upper schemes, and S2a–c). If the monkeys recognized the value during the initial stimulus presentation and anticipated it before the second finger insertion, this guided their value-based finger insertion behavior. In an example session, the monkey demonstrated quicker finger insertions in response to the second finger-in cue when anticipating good stimuli than when anticipating bad stimuli in both tasks (two-tailed unpaired t-test, p = 2.646 × 10^−6^ in T-VRT; p = 2.116 × 10^−6^ in V-VRT) (Fig. 2e).

We analyzed the differences in finger insertion RTs across all sessions in both tasks. In the T-VRT, 84% of sessions (255 out of 299) and in the V-VRT, 91% of sessions (273 out of 299) showed significant differences in finger insertion RTs, with faster finger insertion for anticipated good stimuli than bad stimuli (p < 0.05, two-tailed unpaired t-test for each neuron) (Figs. 2f and S2d–e). In addition, we found that the monkeys were more inclined to avoid touching the bad stimuli than the good stimuli (paired t-test, p = 5.339 × 10^−40^ for T-VRT, p = 1.984 × 10^−70^ for V-VRT) (Fig. S2f). The acquisition of tactile and visual values was further confirmed by their preference for good stimuli in choice trials (Fig. 2c and d, lower schemes). The monkeys successfully selected good stimuli more than bad stimuli in both tasks (paired t-test, p = 2.655 × 10^−215^ for T-VRT, p= 6.758 × 10^−312^ for V-VRT; 92.72 ± 0.48% and 97.66 ± 0.25% average correct choice rate for the T-VRT and V-VRT, respectively) (Figs. 2g, S2g, and S2h) (Movies S2). This value-guided finger insertion behavior indicates that the monkeys successfully acquired values associated with the tactile and visual stimuli, enabling us to investigate the neural encoding of these tactile and visual values.

### Encoding of both tactile and visual values by individual neurons in the monkey putamen

To examine whether the putamen neurons processed value information, we recorded the spike activity of single neurons in the putamen during these tactile and visual value tasks. In the T-VRT, the monkeys were instructed to insert their index fingers into a hole upon the appearance of the first finger-in cue (Fig. 2c, upper scheme). Following the insertion of the finger, one of the tactile stimuli was presented for a duration of 500 ms to examine the neural responses encoding the value associated with each braille pattern (Figs. 2c and S2a). In the V-VRT, the monkeys were required to insert their index fingers into the hole to display a fractal image on the screen, ensuring consistent motor movements between both tasks for a fair comparison of the neural responses encoding tactile and visual values (Fig. 2d).

We identified value discrimination responses in the putamen neurons. Figure 3a shows an example neuron in the T-VRT that encoded value information during tactile stimulus presentation, with stronger responses to good tactile stimuli than bad ones (referred to as a *stimulus–response value neuron*). Notably, this same neuron also exhibited significantly greater responses to good visual stimuli than bad ones in the V-VRT (Fig. 3b). This example neuron demonstrates that both tactile and visual values were encoded in a single neuron of the putamen.

**Figure 3.**
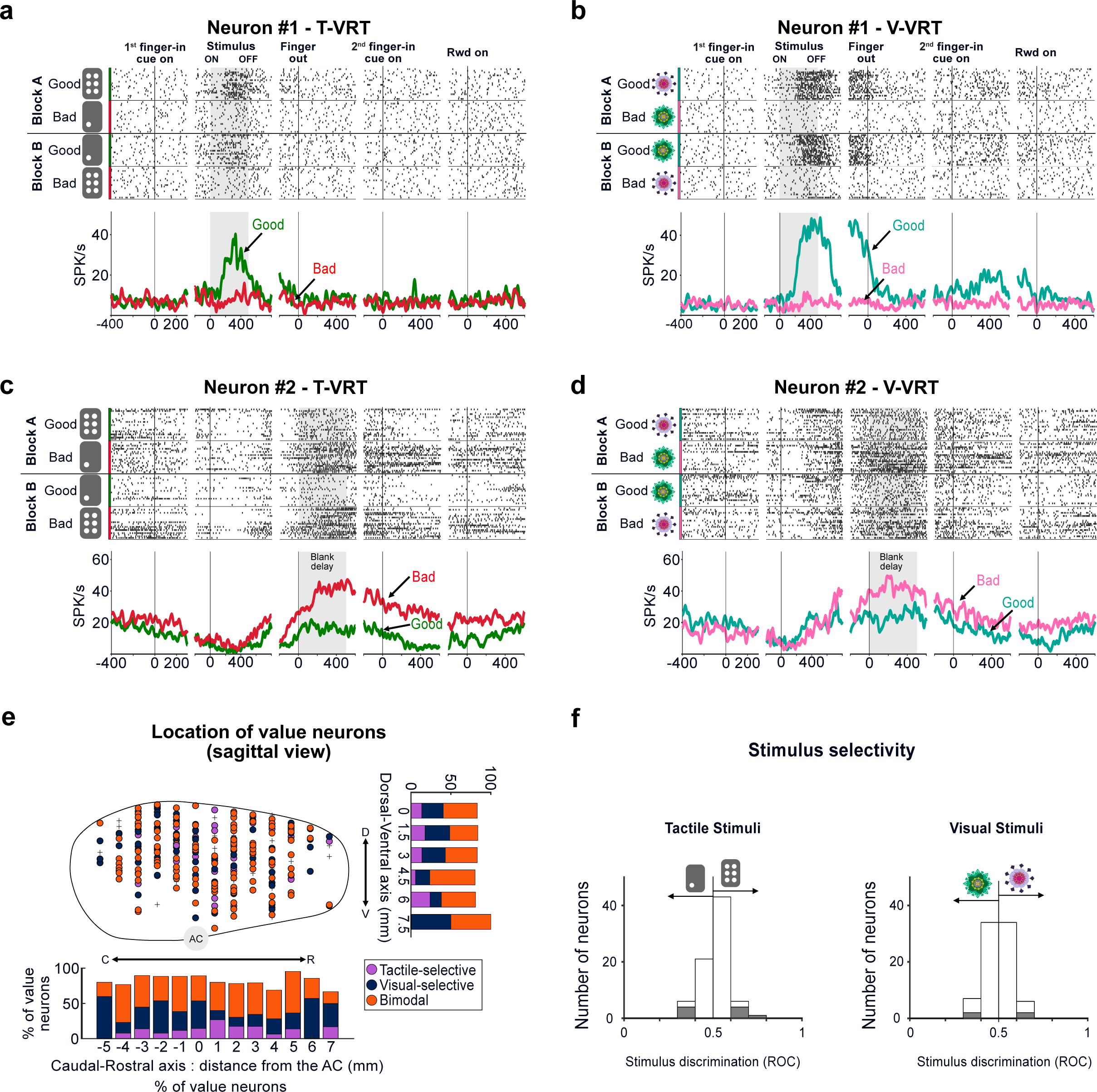
Both tactile and visual values encoded in single neurons of the monkey putamen. **(a–b)** Example of an individual putamen neuron encoding both tactile and visual values during the stimulus presentation period. Neuronal responses to good and bad stimuli are shown in raster plots and peristimulus time histograms (PSTHs). **(c–d)** Example of a single putamen neuron encoding both tactile and visual values during the blank delay period. Responses of this neuron in the same format as in (a–b). **(e)** Locations of value neurons in the sagittal view. Purple and navy indicate tactile-(nL=L41) and visual-selective (nL=L77) value neurons, respectively. Orange indicates bimodal value neurons (nL= 129). Black crosses indicate non-value neurons (nL=L52). D, dorsal; V, ventral; R, rostral; C, caudal; AC, anterior commissure. **(f)** Stimulus discrimination responses of value neurons. For each neuron, the difference in responses to two stimuli was calculated as the ROC area. The gray bars indicate neurons that exhibited statistically significant stimulus selectivity responses in the T-VRT (n = 77) and V-VRT (n = 81).

Furthermore, we discovered another type of neuron that encoded value information during the blank delay period when the monkeys remained nearly motionless after retracting their fingers (referred to as a *delay value neuron*) (Fig. 2c and d). The example delay value neuron in Figure 3c exhibited a higher response to bad tactile stimuli than good tactile stimuli during the delay period but not during stimulus presentation in the T-VRT. In the V-VRT, this same neuron also showed value discrimination responses for visual stimuli during the delay period (Fig. 3d). Our example neural data thus reveal that the putamen contains individual stimulus–response and delay value neurons that encode both tactile and visual value information (referred to as *bimodal value neurons*).

### Three types of value neurons in the primate putamen

We discovered these bimodal value neurons distributed across the putamen, but we also found modality-selective value neurons that exclusively encoded either tactile or visual values (Figs. 3e and S3a). Among the 299 task-responsive neurons found in the putamen, 247 (82.6%) showed value discrimination responses to both tactile and visual stimuli or to either one. To confirm that these value discrimination activities were not influenced by different finger movements in response to good and bad stimuli, we analyzed the movement patterns during the first finger insertion period (Fig. S4a). Our analysis showed no differences in finger movements when monkeys interacted with stimuli of different values, indicating that the value discrimination responses were not a result of different finger movements (Fig. S4b– c).

The average responses of these three types of value neurons showed distinct value discrimination activities (Figs. 4a–f). Considering that various putamen neurons responded more strongly to either good or bad stimuli (*positive* and *negative value-coding neurons* in Fig. S5), we calculated the average responses based on the neurons’ preferred and non-preferred values. The population activities of the stimulus–response value neurons increased after the presentation of the first finger-in cue, and they displayed distinct responses encoding tactile values, visual values, or both (Fig. 4a–c). *Tactile-selective value neurons* responded more strongly to the preferred tactile value than the non-preferred one but did not show value discrimination responses to visual stimuli (Fig. 4a). In contrast, *visual-selective value neurons* showed stronger responses to their preferred visual value than the non-preferred one, but they did not exhibit value discrimination activity to tactile stimuli (Fig. 4b). Different from these modality-selective value neurons, bimodal value neurons showed value discrimination responses to both tactile and visual stimuli during stimulus presentation (Fig. 4c).

**Figure 4.**
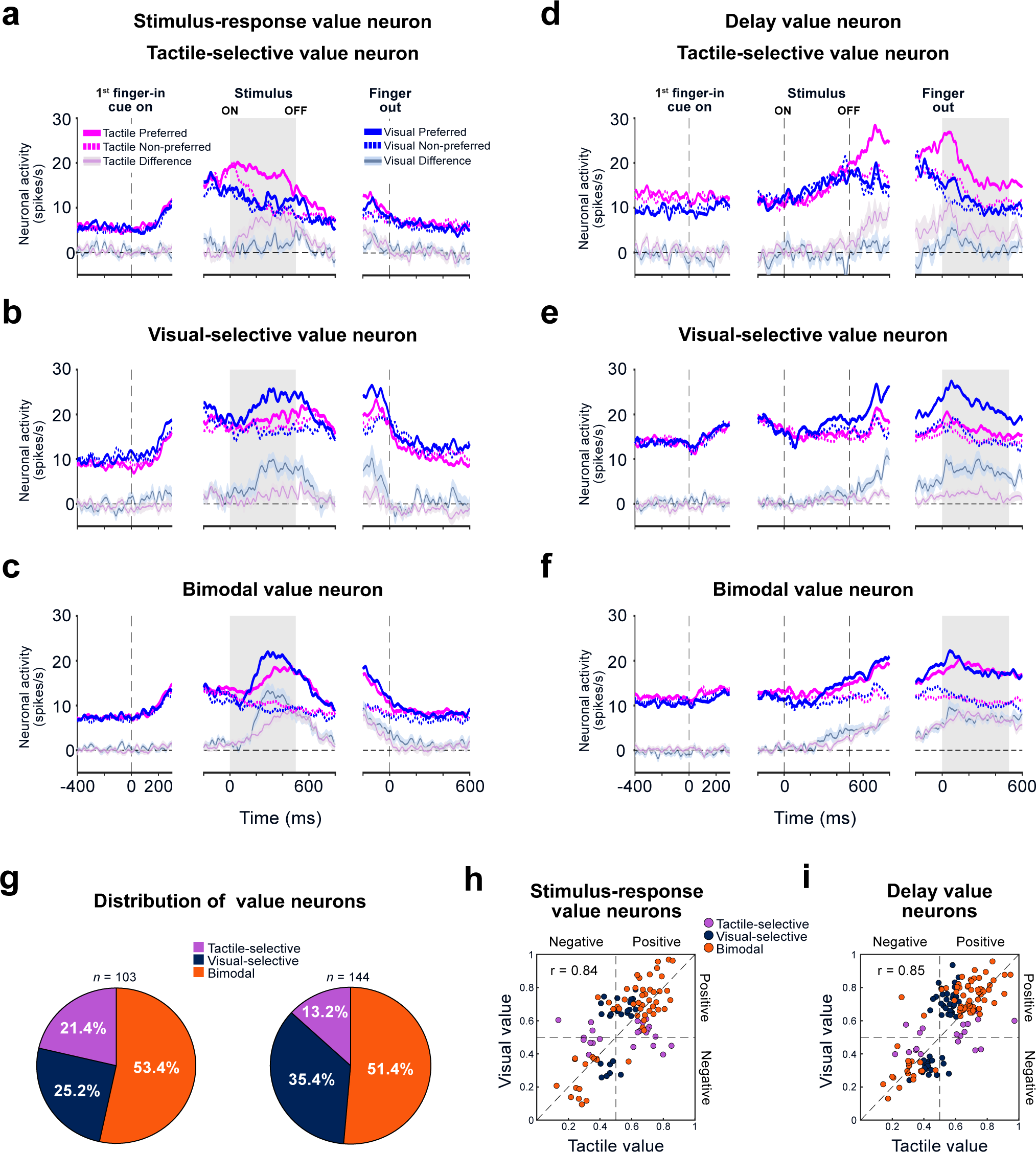
Half encoded both values, and the other half encoded modality-selective values. **(a–c)** The population responses of each type of value neuron in the putamen during the stimulus presentation period. The PSTHs indicate the averaged activities of tactile-selective (n = 22) **(a)**, visual-selective (n = 26) **(b)**, and bimodal (n = 55) **(c)** value neurons to good and bad stimuli during the T-VRT and V-VRT. The value discrimination activities in each task are indicated by transparent lines (mean ± SE). **(d–f)** Population responses of each type of value neuron in the putamen during the blank delay period. The same format as in (a–c) for tactile-selective (n = 19) **(d)**, visual-selective (n = 51) **(e)** selective, and bimodal (n = 74) **(f)** value neurons. **(g)** Distribution of the three types of value neurons in the putamen. The left and right pie charts show the distributions of stimulus–response and delay value neurons, respectively. **(h)** Comparison between the tactile value coding (abscissa) and visual value coding (ordinate) of stimulus–response neurons in the putamen (n = 103). For each neuron (each dot), the magnitudes of the tactile and visual value coding, calculated by ROC areas, are plotted. ROC areas higher and lower than 0.5 indicate positive and negative value coding, respectively. **(i)** Comparison between tactile value coding and visual value coding of delay neurons in the putamen (n = 144). The format is the same as in (h).

Similar patterns of value discrimination activity were found in the averaged responses of the delay value neurons after the stimulus disappeared. They showed three types of value-coding responses during the blank delay period after the finger was retracted (Fig. 4d–f).

### Electrophysiological features of putamen value neurons

The spike shape and duration of these value neurons were not different from those of the non-value neurons, suggesting that these value neurons were the medium spiny neurons typically found in the putamen (one-way ANOVA, F(3, 409) = 2.35, p = 0.071 for spike duration) (Fig. S3b and c). In the comparison of baseline activities, visual-selective value neurons exhibited slightly higher activity levels than tactile-selective and bimodal value neurons, suggesting a potential difference in their input pathways (one-way ANOVA, F(3,594) = 3.94, p = 0.008; post-hoc Bonferroni pairwise comparison, p = 0.012 for tactile-selective and visual-selective; p = 0.037 for visual-selective and bimodal) (Fig. S3d).

We next analyzed the stimulus selectivity of these value neurons by comparing their neural responses to pairs of tactile or visual stimuli associated with reward. 11.68% (9/77) of value neurons showed stimulus selectivity to particular tactile stimuli, and 3.93% (4/81) exhibited selectivity to specific visual stimuli during the initial stimulus presentation (two-tailed Wilcoxon rank-sum test, p < 0.05) (Fig. 3f). The data suggest that these value neurons, which are widely distributed in the putamen, are primarily involved in processing value information rather than stimulus selectivity.

### Half of value neurons converge tactile and visual value information

Interestingly, more than half of the value neurons (52.23%, 129 out of 247) were bimodal value neurons that converged both tactile and visual values in single neurons. These proportions were consistently observed across both monkeys (51.68% and 53.06% in UL and EV, respectively). 53.4% (55 out of 103) of stimulus–response value neurons and 51.39% (74 out of 144) of delay value neurons were bimodal value neurons, indicating their consistent distribution across these categories of value neurons (Fig. 4g).

Their value discrimination responses to tactile stimuli showed a strong positive correlation with the discrimination responses to visual stimuli in both bimodal stimulus– response and delay neurons (r = 0.84, p = 6.247 × 10^−16^ and r = 0.85, p = 1.148 × 10^−22^, for stimulus–response and delay neurons, respectively, Pearson correlation coefficient) (Fig. 4h and i). 95.35% of bimodal value neurons encoded tactile and visual values in the same direction, either positive or negative, with only 4.65% of these neurons encoding the values in opposite directions. These results suggest that each bimodal value neuron processes value information in a similar manner in terms of value-encoding direction and strength, regardless of the sensory modality presented.

In the other half of value neurons, 21.36% (22 out of 103) of stimulus–response value neurons and 13.19% (19 out of 144) of delay value neurons exhibited neural responses selective to tactile value (Fig. 4g). The remainder of these value neurons, 25.24% (26 out of 103) for stimulus–response neurons and 35.42% (51 out of 144) for delay value neurons, selectively encoded visual values (Fig. 4g). These findings indicate that the putamen processes value information through a combination of parallel and convergent information pathways at the single-neuron level, with more than half of neurons converging tactile and visual value information.

### The responses of value neurons correlated with anticipated value-guided finger insertion

We have demonstrated that the putamen neurons encoded values independently of movements during the first stimulus presentation (Figs. 4 and S4). These value discrimination activities might influence subsequent behavior, resulting in an observable correlation between neural activity and the second finger insertion (Fig. 2c and d). To examine how the neural activity changed as finger insertion behavior adapted to value reversals, we analyzed the correlation between RTs for the second finger insertion and the activities of value neurons during stimulus presentation and the delay period (Figs. 5a–b, and S6).

**Figure 5.**
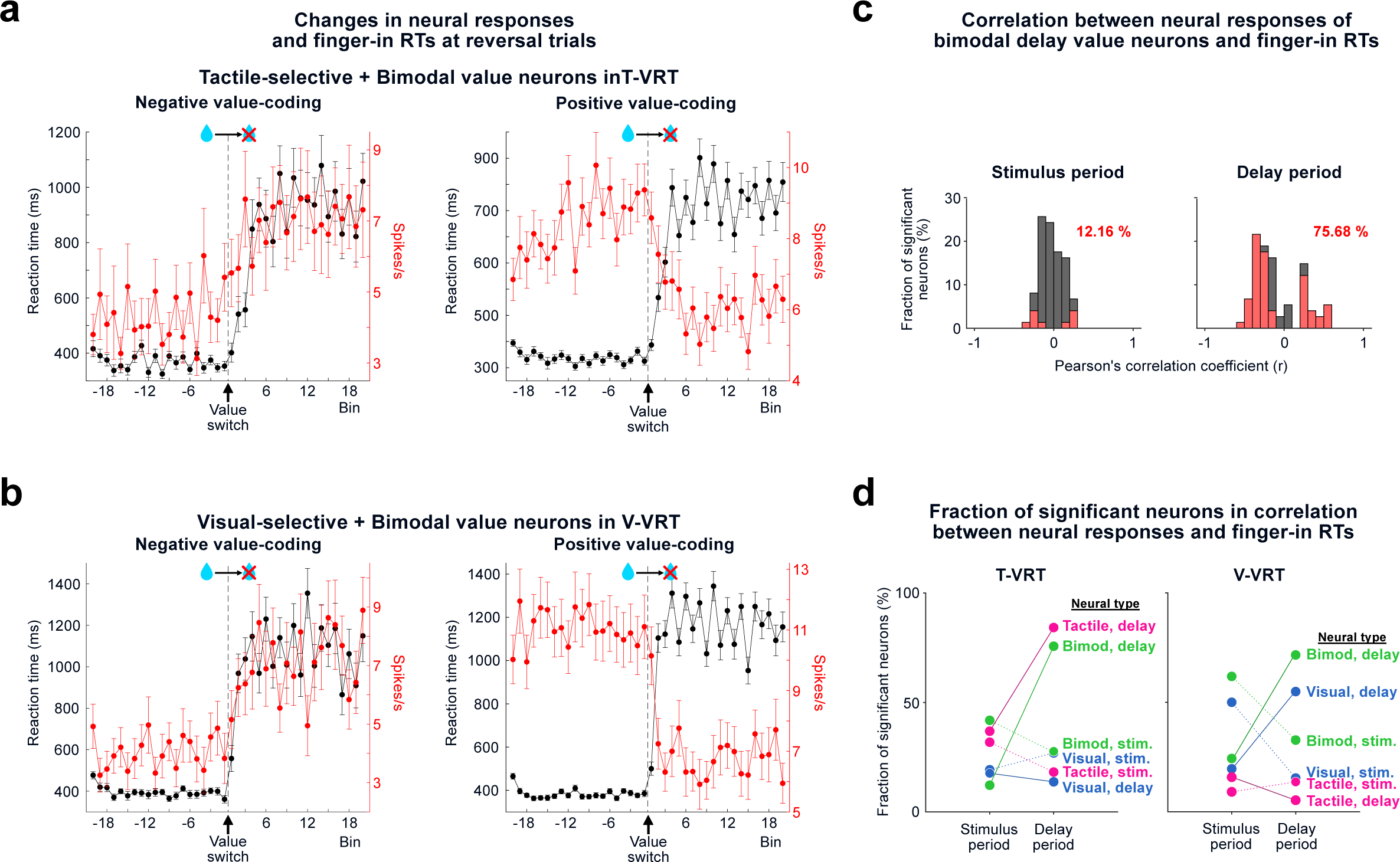
Neural responses correlated with anticipated value-guided finger insertion. **(a)** Changes in neural responses and finger insertion speeds across trials in the T-VRT. The averaged spike rates of tactile-selective and bimodal value neurons (red) and reaction times of finger insertion in response to the second finger-in cue (black) are plotted before and after the value switching trials (mean ± SE). **(b)** Plot with the same format as in (a) for visual-selective and bimodal value neurons in the V-VRT. **(c)** Example correlation coefficient histograms of bimodal delay value neurons in the T-VRT (n = 74). The correlation coefficient values were calculated by comparing between the reaction times of finger insertion and neural responses during the stimulus presentation period (left) and blank delay period (right). Red bars indicate neurons with statistically significant correlations (Pearson correlation coefficient, p < 0.05). The percentage indicates the fraction of neurons showing statistically significance in correlation. **(d)** Dynamic changes in the fractions of value neurons correlated with anticipated value-guided finger insertion. The fractions of 6 types of value neurons showing significant correlation coefficients during the stimulus and delay periods are plotted separately for the T-VRT (right) and V-VRT (left).

When the value of each stimulus shifted from good to bad, the speed of finger insertion gradually decreased, possibly as the monkeys recognized the change in stimulus value (from 370.18 ± 1.77 ms for good stimuli to 996.22 ± 8.87 ms for bad stimuli in the T-VRT and V-VRT, mean ± SE) (Fig. 5a and b). This alteration in behavior was accompanied by a dynamic increase in the neural responses of negative value–coding neurons (Fig. 5a and b, left panels). In contrast, the neuronal responses of positive value–coding neurons decreased as the monkeys began to insert their fingers more slowly after the value shifted from good to bad (Fig. 5a and b, right panels). Similar correlations were observed when the value shifted from bad to good (Fig. S6). These results show that the change in finger insertion speed following the value switch coincided with changes in the neural responses of value neurons, suggesting their potential role in guiding finger movements driven by anticipated value.

We discovered two types of value neurons classified by when they encoded values: stimulus–response and delay value neurons (Fig. 4). Considering that the responses of the delay value neurons aligned with the second finger movements more closely than the other type, it raises the question of whether they play distinct roles in guiding anticipated value– driven finger movements. To determine this, we calculated the correlation between the neural activities of each type of value neuron and individual finger-in RTs. Figure 5c shows an example result: 75.68% of bimodal delay value neurons exhibited statistically significant correlations between their responses during the delay period and the RTs for the second finger insertion (p < 0.05, Pearson correlation coefficient). In contrast, only 12.16% of these neurons showed correlations between their responses during the stimulus period and the RTs.

All types of value neurons were analyzed to calculate their correlations with finger-in RTs (Fig. S7). In the T-VRT, the proportions of statistically significant correlations between tactile and bimodal delay value neurons and RTs during the delay period selectively increased, compared with the stimulus period (from 36.84% to 84.21% for tactile selective; from 12.16% to 75.68% for bimodal) (Fig. 5d, left panel). Similarly, in the V-VRT, the proportions of visual and bimodal delay value neurons selectively increased during the delay period compared with the stimulus period (from 19.61% to 54.9% for visual selective; from 24.32% to 71.62% for bimodal) (Fig. 5d, right panel). These findings suggest that delay value neurons played a predominant, modality-selective role in guiding the finger insertion behavior driven by anticipated values.

### Efficient processing of value information by bimodal value neurons

Our finding of neurons that integrate tactile and visual value information leads to a fundamental question: What advantages does convergent information processing offer for encoding values? One potential advantage is requiring fewer neurons to encode values, compared with parallel processing. Because bimodal value neurons can participate in generating both tactile and visual value discrimination responses, the total number of value neurons decreases as the number of bimodal value neurons increases (Fig. 6a). Importantly, the number of neurons dedicated to processing each value remains consistent with that in parallel processing.

**Figure 6.**
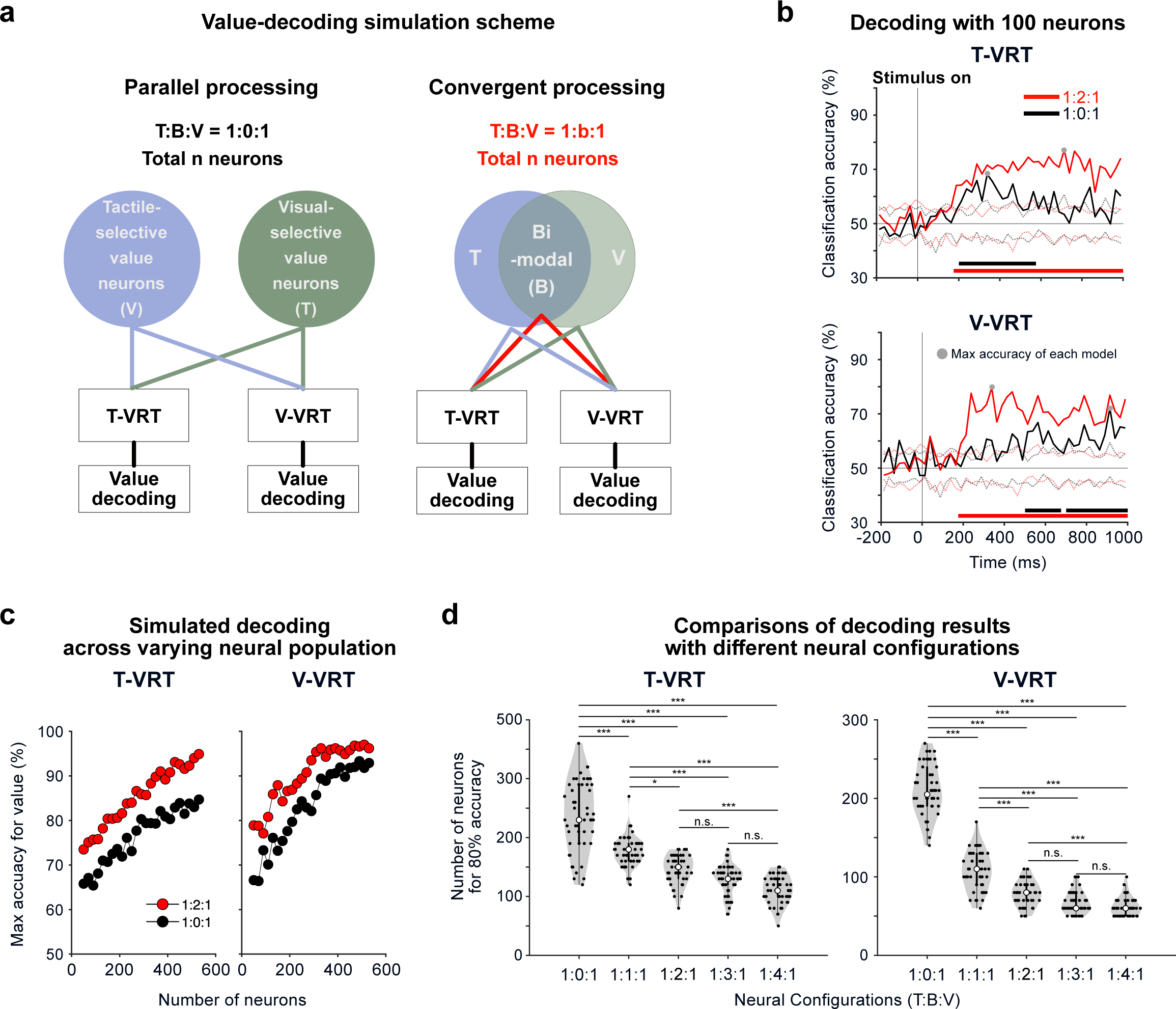
Simulation of parallel and convergent value processes using modality-selective and bimodal value neurons. **(a)** Simulation scheme for value decoding with various neural configurations representing the parallel and convergent processes. Value decoding accuracies were calculated with neural population activities with bimodal value neurons (convergent process, right) and without them (parallel process, left). **(b)** Examples of value decoding in parallel and convergent processes. Value decoding accuracies in the parallel value process (1:0:1 ratio of tactile, bimodal, visual value neurons; n = 100) and convergent value process (1:2:1; n = 100) were compared in the T-VRT and V-VRT. The black and red dashed lines indicate the decoding accuracies of the permutation test. Bars represent periods of decoding accuracy above the permutation results. **(c)** Example plots showing the maximum values of decoding accuracy calculated with various total numbers of value neurons in the T-VRT (left) and V-VRT (right). **(d)** Comparisons of the minimum number of neurons needed to achieve an 80% decoding accuracy among 5 possible neural configurations in the T-VRT (left) and V-VRT (right). One-way ANOVA was applied for comparison (∗p<0.05, ∗∗∗p<0.005, post-hoc Bonferroni pairwise comparison).

We conducted simulations using a population decoding analysis to predict value information across various configurations of value neurons^25^. The value-decoding accuracies were first compared between two models, each comprising 100 neurons: the parallel processing model (1:0:1 for tactile selective, bimodal, and visual selective neurons) and the convergent processing model (1:2:1) (Fig. 6b). Notably, value classification accuracies were higher in the convergent processing model than the parallel processing model in both tasks (Fig. 6b). Next, we calculated the maximum accuracies as the count of value neurons increased in these two models (Fig. 6c). Our simulations consistently revealed that the 1:2:1 configuration of convergent processing outperformed parallel processing, achieving a higher accuracy with a lower number of value neurons (Fig. 6c).

To investigate the efficiency of parallel processing in encoding values as the number of bimodal value neurons increases, we calculated the number of neurons needed to achieve an 80% value-decoding accuracy across different configurations of value neurons. Those simulation results revealed that an increase in the proportion of bimodal value neurons significantly reduced the total number of value neurons required to achieve an 80% value-decoding accuracy (one-way ANOVA and post-hoc Bonferroni pairwise comparison, p < 0.05) (Fig. 6d). Our data suggest that as value convergence increases, the efficiency of value encoding improves. However, we did not find significant differences in modality-decoding accuracies between convergent and parallel processing (one-way ANOVA) (Fig. S8).

Overall, our simulations demonstrate that the use of convergent neural processing with bimodal value neurons enables the putamen to encode value information with fewer neurons than would be required for parallel processing while preserving modality information in neural response patterns.

## Discussion

Our fMRI study demonstrated that both tactile and visual values were represented in the human putamen. In our single-unit recording study with monkeys, we discovered that more than half of individual putamen neurons encoded both tactile and visual values, with the other half selectively encoding either value. Notably, our findings indicate that values originating from different sensory modalities, which might be initially processed in cortical areas, converge within single neurons in the putamen. We further addressed the question of whether bimodal value coding neurons provide any advantage in value processing. Neural decoding simulations revealed that this convergent value process requires fewer neurons than the parallel value process and doesn’t lose modality information. These findings suggest that the convergent pathway through the primate putamen facilitates the processing and generation of abstract value information by efficiently using neurons.

### Tactile value processing in putamen neurons for decision-making via the fingertip

Our study revealed that neurons in the primate putamen encoded the values associated with tactile stimuli experienced at the fingertip. Traditionally recognized for its role in motor control and action value processing, the putamen’s role in processing object value has received relatively little attention^26–28^. A recent study demonstrated that reversible inactivation of the marmoset putamen impaired the performance of a visual value reversal task, suggesting its role in object value processing^18^. Our finding highlights a previously uncharted role for the putamen, encoding tactile values, which enables primates to make optimal decisions by collecting information via their fingertips.

### Integration of tactile and visual values in putamen neurons

How do individual neurons in the putamen process both tactile and visual value information? Various funneling models have previously been suggested and debated, and our findings provide insight into how neurons converge these distinct values^9,29^. Neurons in the putamen receive diverse sensory inputs, including tactile and visual inputs, directly from cortical areas^20,30–32^. Furthermore, dopamine inputs from the substantia nigra pars compacta strongly innervate the putamen, enabling it to encode values from various sensory modalities^33^.

These anatomical studies suggest that a potential circuit of axon terminals from individual neurons in the cortex that are responsible for processing either tactile or visual information establishes connections with single neurons in the putamen. Dopamine projection to the putamen might modulate the synaptic strength of these convergent inputs during value learning. This neural pathway allows for the integration of value information from both sensory modalities in single neurons of the putamen. In contrast, putamen neurons encoding either tactile or visual value might receive a single input from the relevant modality-processing area in the cortex.

Overall, our results indicate that both parallel processing and convergent processing occur in the putamen. This raises the important future question of whether the successive structures, such as the globus pallidus and substantia nigra, handle these values through the convergent funneling process.

### Advantages and disadvantages of convergent value processing through bimodal value neurons

One important question is how the brain efficiently processes vast amounts of information with a limited number of neurons. Our findings provide empirical evidence that convergent processing with bimodal value neurons requires fewer neurons to represent values than parallel processing. This, in turn, might conserve resources and increase the brain’s capacity for processing other types of information.

On the other hand, this convergent process seems to retain only value information, omitting sensory modality–specific details, implying that each bimodal value neuron encodes an abstract form of value. Consequently, it becomes challenging to discriminate which modality is processed based on the averaged responses of bimodal value neurons (Fig. 4c and f). However, we observed that the neural response patterns preserved this modality information, as shown in the decoding analysis (Fig. S8). This finding suggests that putamen neurons might use their population response patterns to preserve the compressed representation of sensory modalities^8^.

### Possible roles of the parallel value process through modality-selective value neurons

Our study also showed that nearly half of putamen neurons selectively process either tactile or visual value, suggesting the presence of parallel processing through the putamen. These modality-selective value neurons could serve three possible functions.

First, modality-selective value neurons could act as markers of sensory modality. In cases in which bimodal value neurons exclusively process both tactile and visual values, it becomes challenging to differentiate the sensory source based solely on population average responses. Each modality-selective value neuron might function as a distinct identifier for its associated sensory modality, simplifying the interpretation of sensory information. Second, these neurons might form selective connections with specific motor output regions. For example, visual- and tactile-selective value neurons in the putamen could project to distinct areas controlling eye and forelimb movements, respectively, in the globus pallidus internal segment^6,34^. These modality-selective circuits could be involved in guiding a particular movement, such as saccades or grabbing. Third, these modality-selective value neurons come into play when conflicting value information arises from two sensory modalities. In such value conflicts, bimodal value neurons might struggle to differentiate between conflicting values from different modalities. Conversely, each type of modality-selective value neuron might independently process its specific value to guide behavior when value conflicts occur.

### Transitioning from value recognition to finger-in action: The possible roles of stimulus– response and delay value neurons in prefrontal-striatal circuits

In our value tasks, we temporally separated the phases for value recognition and finger-in preparation. This allowed us to identify two populations of neurons: stimulus– response and delay value neurons. Notably, we found a small number of neurons that sustained their value discrimination responses from the stimulus presentation period to the delay period (10.12%, 25 out of 247 value-coding neurons). Most value-coding neurons exhibited a stronger preference for encoding values during either the stimulus presentation or delay period, indicating their predominant involvement in one of these periods (Fig. 4a–f).

Interestingly, previous studies have reported the presence of two types of neurons in the prefrontal cortex (PFC) that showed stronger responses during either cue presentation or the delay period^35–38^. Considering these two types of PFC neurons, it is conceivable that the two distinct types of value neurons in the putamen are connected to the PFC through different pathways to serve distinct functions. Value information might first be processed by stimulus– response value neurons and then subsequently transferred to the PFC, where it can be modified through interaction with other information. Then, the modified value information could be transmitted to delay value neurons in the putamen to guide finger-in actions using that value information.

## ACKNOWLEDGMENTS

This work was supported by the Neurological Disorder Research Program (NRF-2020M3E5D9079908) through the National Research Foundation (NRF) of Korea, and Korean government (MSIT) grant (NRF-2019R1A2C2005213). We thank D.I. Ko for technical support and members in ILAR, SNU for discussion and technical assistance.

## Author contributions

S.-H.L. and H.F.K. designed and supervised the entire project. SH. H., D. P., JW. L., S.-H.L., and H.F.K. performed the behavior, fMRI, and single-unit recording experiments. SH. H., D. P., and JW. L. analyzed the data and prepared the figures. SH. H., D. P., JW. L. wrote the first draft, and SH. H., D. P., JW. L., S.-H.L., and H.F.K. interpreted data and wrote the final manuscript.

## STAR Methods

### I. MRI study with human participants

#### Participants

Twenty-two neurologically intact right-handed participants (mean age 24.36 ± 3.58 years, range 20–33, 10 females) took part in the experiment. The participants reported that they had normal or corrected-to-normal vision. All participants provided informed consent for the procedure. The experimental procedure was approved by the Institutional Review Boards of Seoul National University and the Korea Advanced Institute of Science and Technology.

#### Stimuli

As tactile stimuli, we used two braille patterns engraved on the surfaces of polyoxymethylene blocks (Fig. 1a). The dimensions of each tactile block were 18 × 18 × 12 mm, and each dot had a diameter and height of 2 × 1 mm. The tactile stimuli were automatically presented via a pneumatic tactile stimulus delivery system, as previously reported^39^. For the visual stimuli, we used two fractal images created with Fractal Geometry (Fig. 1a)^40,41^. The mean luminance across the images was equalized using the Spectrum, Histogram, and Intensity Normalization and Equalization (SHINE) toolbox written with MATLAB (www.mapageweb.umontreal.ca/gosselif/shine). Grayscale images of the fractal objects were used to minimize the color effect on brain responses. The fractal images were presented with a visual angle size of 7 x 7 degrees.

#### Task design: Value reversal tasks for human participants

Participants performed two distinct value reversal tasks, the tactile task and the visual task, while their neural activities were monitored using fMRI. Each task was executed in separate runs, with participants being informed about the upcoming task before each run. In both tasks, tactile and visual stimuli were presented concurrently. Specifically, the tactile task exclusively linked tactile stimuli, such as the braille T-a and T-b, with monetary values, with one stimulus predicting monetary gain (good), and the other predicting monetary loss (bad) (Fig. 1b). The visual stimuli in the tactile task were unrelated to the monetary outcome. Conversely, in the visual task, the fractal images were associated with monetary values. One image was linked to monetary gain (good), and the other was associated with monetary loss (bad), and the braille stimuli were not linked to the outcomes (Fig. 1b). The rule linking each stimulus to its corresponding monetary value was switched in the middle of each run in both tasks. Each task comprised four scan runs, with trial counts of 16, 18, 18, or 20 for each run. The order of the runs was pseudo-randomized for each participant. The variation in trial numbers within each run was intended to mitigate participants’ ability to predict the occurrence of rule switching. The tactile task and visual task alternated, and the task modality of the first run was counterbalanced across participants.

In each trial of both tasks, the onset of the trial was indicated by the presentation of a blue fixation cross. After 500 ms, the cross changed to red as the “insert finger” signal (Fig. 1c). Participants were instructed to insert their right index finger into the entrance hole and touch a braille stimulus in the hole when the red cross was displayed and to withdraw their finger from the hole when the red color of the fixation cross returned blue (“withdraw finger” signal) 1 s later (Fig. 1c). Participants were concurrently exposed to visual stimuli while experiencing the tactile stimuli. To ensure equal exposure to both tactile and visual stimuli, the visual stimulus was presented only while the finger was in the entrance hole. After a 5 s-interval, participants were asked to decide whether to take the object, contingent upon its potential value derived from feedback in preceding trials or through conjecture. To mitigate the potential interference of pre-planned motor signals with value representation, the plus (indicating ‘take’) and minus (indicating ‘do not take’) buttons were randomly assigned to the upper and lower positions (Fig. 1c). Participants were instructed to promptly press the upper button with their left index finger and the lower button with their left middle finger. The response window lasted for 2 s, followed by a 0.5 s feedback period and a variable inter-trial-interval ranging from 2.5 to 13 s.

As feedback, the monetary gains or losses associated with each trial’s response were displayed just above the fixation cross (Fig. 1c). Before the tasks, the participants were informed that the presented feedback would be added to their compensation after the experiment. When the presented stimulus was associated with monetary gain, selecting the plus button resulted in a gain of 150 won, and selecting the minus button yielded a gain of 50 won. If the presented stimulus was linked to monetary loss, choosing the minus button led to a loss of 50 won, and selecting the plus button resulted in a loss of 150 won. Participants were familiarized with these responses and feedback rules before the task.

Before undertaking the two value reversal tasks, participants underwent a preparatory session inside the scanner to familiarize themselves with the stimuli and the finger movement protocol. This familiarization session encompassed four tactile blocks and four visual blocks with four braille patterns and four fractal images, inclusive of the stimuli used in the subsequent value reversal tasks. The participants performed alternating tactile and visual blocks. During the tactile block, the participants were instructed to insert their right index finger into the entrance hole to touch and discern the given tactile stimulus while they fixated on the center of the screen. During the visual block, the participants were asked to view each fractal image displayed at the center of the screen while tactually exploring a plain surface with their right index finger. Throughout the familiarization session, each stimulus was presented ten times, including one-back response trials (20% of the total trials), wherein participants were required to determine whether the tactile (or visual) stimulus of the current trial matched that of the previous trial. Additional details on this familiarization protocol are available in our previous work^39^.

#### fMRI acquisition

Participants underwent scanning on a 3T Siemens MAGNETOM Trio located at Seoul National University. Echo-planar imaging (EPI) data were acquired using a 32-channel head coil with an in-plane resolution of 2.8 mm x 2.8 mm and 40 2.5 mm slices (0.25 mm inter-slice gap). The scanning parameters were a repetition time (TR) of 2000 ms, echo time (TE) of 25 ms, matrix size of 76 x 76, and field of view (FOV) of 210 mm. Whole brain volumes were scanned, and slices were oriented approximately parallel to the base of the temporal lobe. Following the experimental runs of the fMRI session, anatomical images were acquired using the standard magnetization-prepared rapid-acquisition gradient echo (MPRAGE) sequence.

#### fMRI analysis

fMRI data were analyzed using AFNI (http://afni.nimh.nih.gov), SUMA (AFNI surface mapper), FreeSurfer, and custom MATLAB scripts. Standard data preprocessing procedures, including slice-time correction and motion correction, were conducted.

To identify regions encoding value information within the human striatum, event-related brain activation patterns in the BOLD signal were initially extracted at the onset of tactile and visual stimulus perception in each trial. This extraction was performed using a standard general linear model in the AFNI software package (3dDeconvolve using the GAM option with motion parameters), and individual trials were modeled as separate events. The estimated t-value for each trial was used in the searchlight analysis^42^. Searchlight spheres, each with a radius of 9.8 mm (corresponding to approximately 123 voxels) were defined within the striatum mask. This mask encompassed the bilateral caudate, putamen, and nucleus accumbens, automatically delineated by the parcellation of FreeSurfer (‘aparc.a2009s+aseg_REN_all.nii.gz’). The t-values extracted from the voxels within each sphere across the striatum were then normalized within each voxel by subtracting the mean value across all stimulus conditions. To derive value discrimination indices, Pearson correlation coefficients were calculated between the voxel patterns of within-value trials (good-and-good or bad-and-bad) and those of between-value trials (good-and-bad). These correlation coefficients underwent Fisher’s z transformation, and the value discrimination indices were defined as the mean within-value correlations subtracted by the mean between-value correlations. When calculating discrimination indices, trials involving incorrect button responses, the first trial of each run, and rule-switching trials were excluded, and all remaining possible pairs of trials from different runs were used. The results were registered to the MNI space from each participant’s original volume for the subsequent group analysis.

#### Statistical analysis

To compare behavioral performance between the tactile and visual tasks, a paired t-test (two-tailed) was conducted in MATLAB. To assess the significance of the value discrimination indices against the baseline level (zero), one-sample t-testing (right-tailed) was used through the AFNI software function (3dttest++).

### II. Electrophysiology study with macaque monkeys

#### Braille presentation for non-human primates

We developed an apparatus for presenting braille patterns to non-human primates (Figs. 2a and S1). The braille presenter comprises pairs of four main components: pneumatic pin cylinders, braille units, photo interrupters as finger sensors, and endoscopic cameras (Figs S1a-b).

Each braille presentation case includes a finger entrance hole to allow tactile exploration of the braille pattern presented inside (Fig. S1b). A braille unit consists of six dots, each independently controlled by an individual pin cylinder placed below the braille unit (Fig. S1a). Each pin cylinder vertically moves a wire connected to a braille dot through air pressure delivered by a compressor (KC-T002, Korea). The computer program BLIP, designed for electrophysiology and behavior studies with macaque monkeys (National Institutes of Health, USA), was customized to control the movement of each braille pin in accordance with the task schedule^39,41^.

To precisely measure the timing of finger insertion into the hole in the braille presentation case, two photo interrupters (F249, Arduino module) were placed on opposite sides of the finger entrance hole (Fig. S1b). To further ensure real-time monitoring of the monkeys’ finger insertion movements and braille pattern presentation, endoscopic cameras (PS-EC200, Korea) were installed at each finger entrance hole (Fig. S1c). We additionally monitored the monkeys’ arm movements during the experiment using a camera (D455, Intel, USA) installed above the braille presenter.

#### Visual stimuli

We used two fractal images created using Fractal Geometry for the V-VRT^40,41^ (Fig. 2b). The mean luminance was equalized across the images using the SHINE toolbox written with MATLAB (www.mapageweb.umontreal.ca/gosselif/shine).

#### Tactile stimuli

Braille patterns were used as the tactile stimuli in the T-VRT. Each braille unit consists of six dots, which were made of an aluminum alloy (Fig. S1b). The diameter and height of each dot was 2 × 1.5 mm, and the size of each braille unit was 10 × 5 mm.

#### General procedures

Two adult monkeys (*Macaca mulatta;* 5.2 kg female monkey EV and 10.7 kg male monkey UL) were used for the non-human primate experiments. Animal care and experimental procedures were approved by the Seoul National University Institutional Animal Care and Use Committee. Under general anesthesia and in surgical conditions, a plastic head holder and a recording chamber were implanted onto the monkeys’ skulls. The position of each chamber was tilted laterally by 25° so that it corresponded to the putamen. We started the training and recording sessions after the monkeys were fully recovered from the surgery.

#### Behavioral tasks

The behavioral tasks were controlled by a Window-based experimentation data acquisition system, BLIP (National Institutes of Health, USA) (www.cocila.net/blip). The monkey sat in a primate chair facing a frontoparallel screen in a sound-attenuated and electrically shielded room. The visual stimuli, squares and fractals, were approximately 20°LxL20° in size. These stimuli were presented on an LCD monitor with a 120-Hz refresh rate (29WL500, LG, Korea). An infrared camera was used to record the monkey’s eye position with a sampling rate of 1 kHz (Bio-signals, USA). The BLIP software precisely monitored the timing of visual stimulus presentation using a signal from a photodiode attached to the monitor. Tactile stimuli in the form of braille patterns were presented using the braille presenter, as described above.

#### Tactile value reversal task and visual value reversal task for macaque monkeys

To test whether individual neurons converged values from different sensory modalities, we designed tasks in which each tactile or visual stimulus could be associated with a liquid reward outcome (Figs. 2c–d and S2b–c). The two tasks using visual (V-VRT) and tactile (T-VRT) modalities were introduced while recording a single neuron. Each task was composed of single-stimulus trials and choice trials.

To examine tactile and visual value responses in individual neurons, we used a value reversal procedure in which the stimulus-value contingency was reversed in each block of 50–60 trials (Fig. 2b). In the first block of T-VRT or V-VRT, one stimulus was associated with a liquid reward, and the other was not. In the subsequent block, the object-reward contingency was reversed. The selection of the stimulus associated with a liquid reward in the initial block was randomized.

#### T-VRT procedure

In the single-stimulus T-VRT trials, the monkeys were trained to insert their left index finger into the left hole of the braille presentation case to experience the braille pattern when the finger-in cue appeared on the screen (colored square cues). To ensure precise exposure to the tactile stimulus during the first stimulus presentation, one of two braille patterns (1-dot and 6-dot) was delivered to the monkeys 50ms after their index fingers entered the hole, and the delivered braille pattern was maintained for 500ms (Fig. S2a). The monkeys were required to maintain contact with the tactile stimulus until the finger-in cue disappeared along with the stimulus. The monkeys had to rely solely on recognizing the value of the tactile stimulus with their tactile perception because the braille pattern presented in the opaque braille presentation case was not visible. After finger withdrawal, a blank black screen was presented for a randomized duration ranging from 500 to 1000ms.

When the second finger-in cue appeared on the screen, the monkeys could insert their finger back into the hole to touch the same tactile stimulus experienced during the first finger insertion. However, the monkeys were free to decide whether to insert their finger or not when the second finger-in cue presented. To receive the reward when the stimulus associated with a liquid reward was presented, the monkeys were required to maintain their finger insertion for 200ms. The reaction time for finger insertion was measured as the time from the presentation of the second ‘finger-in’ cue to the actual finger insertion (Fig. S2a).

A choice trial followed every four single-stimulus trials. In the T-VRT choice trial, two square cues for choice were presented on the screen, and two different braille patterns were simultaneously displayed in the two holes in the braille presentation case. The monkeys were trained to freely explore these two braille patterns by inserting their left fingers into each hole. To choose one stimulus, monkeys had to maintain their finger insertion for more than 1500ms. The reward was provided after the monkeys completed the choice finger insertion (Figs. 2c–d and S2b–c, lower scheme). For monkey UL, the choice square cues were simply presented at the beginning of the trials (Fig. 2c–d). For monkey EV, however, the choice trial commenced with the first ‘finger-in’ cue instead of two choice cues. EV was required to insert a finger into the hole for 500ms. After finger withdrawal and the blank delay period, two square cues for choice were presented on the monitor (Fig. S2b–c).

#### V-VRT procedure

The procedure for the V-VRT was the same as for the T-VRT, except for the presentation of visual stimuli instead of tactile stimuli (Figs. 2c–d and S2b–c). To ensure consistent finger movements in both the T-VRT and V-VRT, the monkeys were also trained to insert their left index finger to display the fractal images in the V-VRT. Additionally, to minimize experimental confounding factors from the braille pattern-raising acoustic noise, a dummy braille pattern in the unused right hole was presented during the single-stimulus trials. This ensured that monkeys experienced a similar level of noise in both the V-VRT and T-VRT.

In the single-stimulus trials, the first finger-in cue was replaced with one of two fractal images by finger insertion. The monkeys were required to keep inserting their finger for 500ms. After finger withdrawal and the blank delay period, the second ‘finger-in’ cue appeared on the screen. The monkeys were allowed to freely decide whether to insert their fingers or not upon the second cue. When the monkeys reinserted their finger into the hole, the same fractal image presented during the first finger-in period was displayed for 200ms. The liquid reward was delivered 200ms after finger reinsertion.

In the choice trials, two square cues for choice were presented on the screen. The monkeys were free to insert their fingers into the holes to choose one fractal image. Each fractal image was presented to the monkey after they inserted their finger into the hole. The monkeys selected a fractal image by maintaining finger insertion for 1500ms. The reward was delivered upon completion of the monkeys’ choices. The overall procedures for the choice trials for each monkey were the same as those in the T-VRT.

#### Single-unit recording

While the monkey performed a task, the activity of single neurons in the putamen was recorded using conventional methods. The recording sites were determined using a 1mm spacing grid system, with the aid of MR images (3T, Siemens) obtained along the direction of the chamber. Single-unit recording was conducted using a glass-coated electrode (Alpha-Omega). The electrode was inserted into the brain through a stainless-steel guide tube and advanced by an oil-driven micromanipulator (MO-974A, Narishige). Neuronal signals from the electrode were amplified, filtered (250 Hz to 10 kHz), and digitized (30-kHz sampling rate and 16-bit A/D resolution) by a Scout system (Ripple Neuro, UT). Neuronal spikes were isolated online using custom voltage-time window-discrimination software (BLIP, Laboratory of Sensorimotor Research, National Eye Institute National Institutes of Health, available at www.cocila.net/blip), with the corresponding timings detected at 1 kHz. The waveforms of individual spikes were collected at 50 kHz.

#### Data analysis Behavioral analysis

To assess the speed of finger insertion into the hole based on anticipated value, we calculated the RT from the presentation of the second ‘finger-in’ cue to the actual finger insertion. The statistical significance of RT differences between good and bad stimuli was validated with a two-tailed unpaired t-test. To illustrate the RT differences across all neural recording sessions, they were calculated by subtracting the averaged RT to good stimuli from the RT to bad stimuli in each session and plotted in a histogram (Figs. 2f and S2d–e).

#### Task-related response

To examine task-related neuronal responses, we isolated the activity of a single neuron and counted the numbers of spikes during the T-VRT and V-VRT. To assess the task-related responses, we counted and compared the numbers of spikes between each targeted test window and control window: control windows were set to 200–0 ms before each task event started, and test windows were set to 0–500 ms after each task event started. The analyzed task events were (1) 1^st^ finger-in cue-on, (2) 1^st^ finger-in, (3) Stimulus-on, (4) Stimulus-off, (5) 1^st^ finger-out, (6) 2^nd^ finger-in cue-on, (7) 2^nd^ finger-in, (8) Reward, (9) 2^nd^ finger-in cue-off, (10) 2^nd^ finger-out. To assess statistical significance in task-related responses, we used the Wilcoxon rank-sum test to compare spike counts between the control and test windows in individual trials.

#### Value-coding activity during the first stimulus presentation

To examine the degree of value discrimination response, we measured the magnitude of each single neuron’s activity in response to each tactile or visual stimulus by counting the numbers of spikes within the test windows in individual trials. To investigate the value response triggered by stimulus presentation, we analyzed neural responses during the stimulus presentation (0–500 ms time window after stimulus onset) in both the T-VRT and V-VRT. The firing rates of individual putamen neurons in response to good and bad stimuli were compared using the Wilcoxon rank-sum test to assess the statistical significance of value discrimination. Neurons exhibiting exclusive value discrimination responses in the T-VRT but not the V-VRT were categorized as tactile-selective value neurons. Conversely, those demonstrating value discrimination responses solely in the V-VRT but not the T-VRT were identified as visual-selective value neurons. Neurons showing value discrimination responses in both the T-VRT and V-VRT were classified as bimodal value neurons.

#### Value-coding activity during the blank delay period

We unexpectedly observed neurons exhibiting stronger value discrimination responses during the blank delay period than during the stimulus presentation period. We thus analyzed neural responses during a 500ms time window following complete finger withdrawal in both the T-VRT and V-VRT. During this time window, the monkeys were presented with only a blank screen to allow us to isolate the value discrimination response from the potential influences of sensory inputs and movements. The same categorization method used for the stimulus–response neurons was applied to classify the types of value neurons during the blank delay period.

Some neurons exclusively exhibited a value discrimination response during the blank delay period or the stimulus presentation period, whereas others showed a more strongly biased response in either period. Neurons were categorized based on their responses in each period using a receiver operating characteristic (ROC) curve calculated with the firing rates of individual neurons to good stimuli versus bad stimuli. A small subset of neurons sustained their value discrimination responses across both periods, leading to the classification of two types of value neurons.

#### Stimulus and value discrimination index

To determine whether value neurons exhibited stimulus-selective responses, we compared spike numbers for each tactile or visual stimulus in individual trials using the Wilcoxon rank-sum test. For an overall illustration of stimulus-selective responses by value-coding neurons, we measured the magnitudes of responses to each braille pattern and fractal image using the area under the ROC curve. Selectivity responses to tactile stimuli were analyzed with spike counts from both tactile-selective and bimodal value neurons. Selectivity responses to visual stimuli were analyzed using spike counts from both visual-selective and bimodal value neurons. An ROC value of 0.5 indicates identical responses to both stimuli, and values closer to 1 or 0 indicate a stronger response to a specific stimulus (Fig. 3f).

We calculated the value discrimination index using an ROC analysis based on the magnitudes of neural responses to good versus bad stimuli in both the T-VRT and V-VRT (Fig. 4h-i). ROC areas higher and lower than 0.5 indicate positive and negative value-coding, respectively

#### Analysis of finger movement during stimulus presentation using DeepLabCut

The movements of the monkeys’ left index fingers were analyzed using DeepLabCut (version 2.2.2) and MATLAB^43,44^. The movements were synchronized and recorded with neural response and behavior data using the Scout system and Trellis software (Ripple Neuro, UT). To detect the positions of the index fingers, three points of the fingernail were manually identified and labeled based on 120 numbers of frames taken from six videos for each monkey (then 95% was used for training) (Fig. S4a). We used a ResNet-50-based neural network with default parameters for 3 number of training iterations. We validated with 1 number of shuffles and found the test error was: 5.33 (for monkey U) or 5.71 (for monkey E) pixels, train: 2.33 (for monkey U) or 3.09 (for monkey E) pixels (image size was 640 by 480). We then used a p-cutoff of 0.6 to condition the X, Y coordinates for consequent analyses. This network was then used to analyze videos from similar experimental settings.

To examine the difference in finger movements between good and bad stimuli, we traced the center of gravity of the fingernail in each frame extracted from the network (Fig. S4a). We computed the Euclidean distances between the centers of gravity in each frame of the trials in the comparison group. These distances were averaged, resulting in a single distance value for each comparison. Differences in the distances indicate that the monkeys exhibited distinct finger movements based on the conditions in the hole. The statistical significance of differences was calculated with a two-tailed unpaired t-test.

#### Correlation between neural response and finger-in reaction time

To examine the correlation between the responses of each neuron and the second finger-in RT across trials, we initially calculated the number of spikes in the respective value-coding windows. To analyze the correlation between the activity of stimulus–response value neurons and finger insertion speeds, spike counting was performed in the 0–500ms window from stimulus onset. To analyze the correlation between the activity of delay value neurons and finger insertion speeds, the analysis included the count of spikes in the 0–500ms after finger withdrawal. To assess how neural responses and RTs changed around the time of value reversal, spike counts in each bin of two trials were averaged and plotted in graphs (Figs. 5a–b and S6).

To analyze dynamic changes in correlations, we calculated the number of spikes for both stimulus–response and delay value neurons in both windows during the stimulus presentation period (0–500ms from stimulus onset) and the blank delay period (0–500ms after finger withdrawal). Subsequently, we investigated whether the second finger-in RTs in each trial correlated with neural counts in these two distinct windows. A Pearson correlation analysis was conducted to assess statistical significance. Based on the coefficient r values, we use a histogram to represent the number of each type of value-coding neurons exhibiting statistically significant correlations (Figs. 5c and S7).

#### Neural population decoding Time window

This analysis was conducted to understand how value information is processed in the population responses of putamen value neurons. In the decoding analysis, both groups of value neurons were considered together using the spike data in two time windows: one during the stimulus period aligned to stimulus onset (−200–500 ms) and the other during the delay period aligned to finger withdrawal (0–500 ms). We calculated the spike train of each 25ms bin within each period and concatenated the spike counts in these two periods for the subsequent decoding analysis.

#### Neural decoding analysis with simulated neural population

To compare the efficiency of parallel and convergent value processes, we performed a neural decoding analysis with modality-selective value neurons (n = 118) and bimodal value neurons (n = 129) derived from our original neural data. This analysis used the maximum correlation coefficient classifier method developed by Meyers^45^. The classifier was trained to discriminate values or modalities of stimuli and calculate decoding accuracies. To simulate various ratios of modality-selective and bimodal value neurons, value neurons from each group were randomly sampled. We used five different ratios, with tactile-selective, bimodal, and visual-selective value neurons distributed 1:0:1 (parallel value process), 1:1:1, 1:2:1, 1:3:1, and 1:4:1 (convergent value processes). To simulate a larger number of neurons than our original dataset, we performed neuron augmentation by creating synthetic neurons through random sampling of actual neuronal responses in trials from each modality group.

In the decoding analysis, the firing rates of individual neurons were collected using a 25ms bin width, as described earlier, and normalized by z-score. Trials from each group were randomly divided into training sets and test sets using 10-fold cross-validation. The classifier was trained on data from 9 splits and tested on the 10th split. This cross-validation procedure was repeated 10 times with a different test split each time. The entire process was replicated over 50 resample runs, creating different random populations of training and test splits in each run. The decoding accuracy data were averaged across these resample runs to obtain a single set of results. This entire procedure was iterated more than 50 times, and the results were used as the main dataset for analysis and visualization.

To estimate the statistical significance of the decoding accuracy, a permutation test was performed by shuffling the labels of each trial variable in each neuron. The same decoding procedure was used to generate a null distribution of decoding accuracies. This procedure was repeated 960 times and generated a fully shuffled decoding null distribution for each time bin. We calculated the time points at which the decoding results were greater than all 960 decoding results of the null distribution. The times when the decoding accuracies significantly exceeded the chance level for at least 5 consecutive bins are indicated as solid bars on the bottom of Fig. 6b.

