## Supplementary figures for "Efficient value encoding through convergence of tactile and visual value information in the primate putamen"

**Supplementary figure 1. Overview of the braille presenting system for non-human primates.**

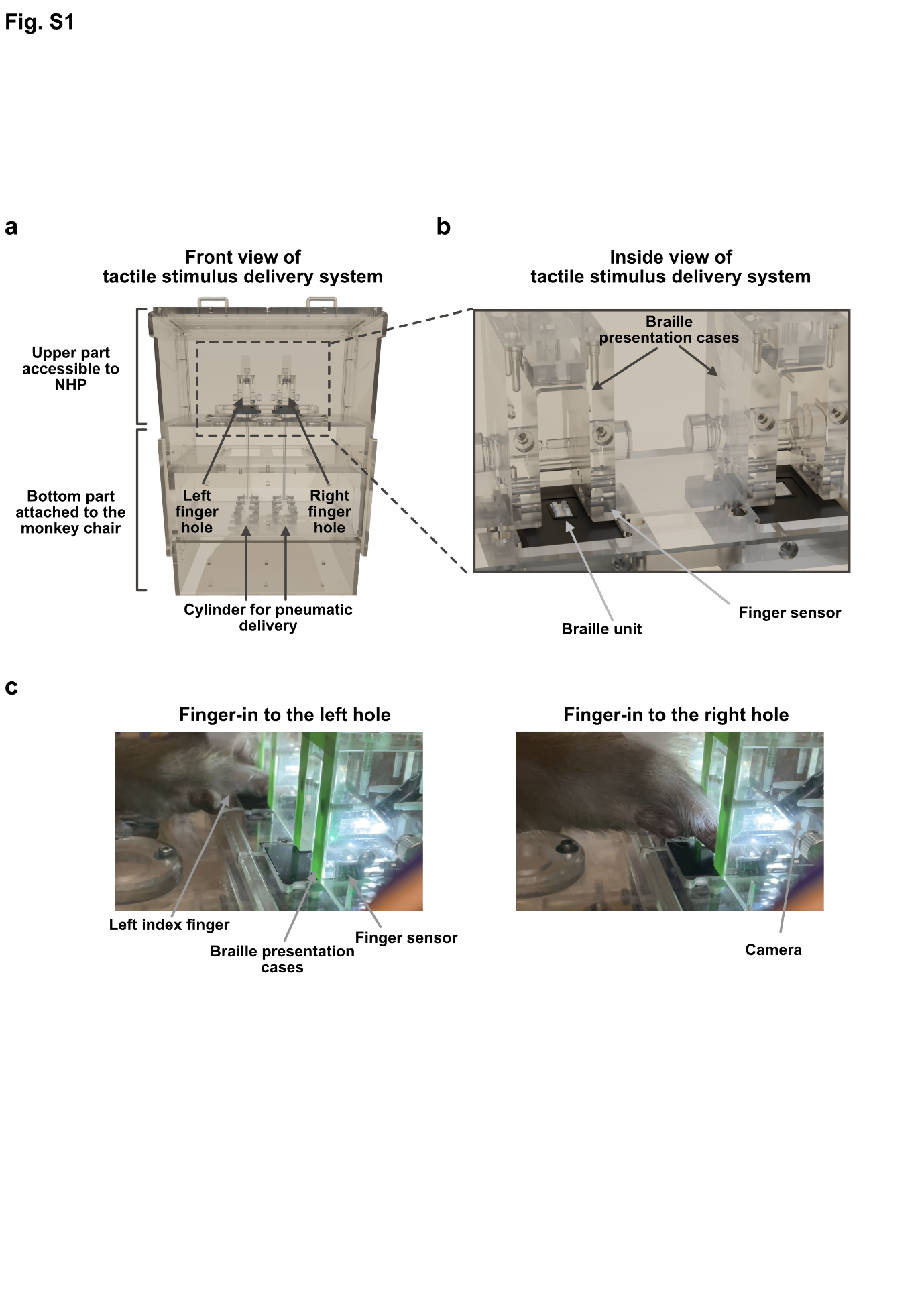
**(a)** A 3-D illustration of the braille presenter in the front view. The bottom part is attached to the monkey chair, and the upper part is accessible to the monkey. **(b)** Magnified view of the braille presentation case. Monkeys inserted their left index fingers into the hole of the braille presentation case. The finger sensor measured the precise timing of finger insertion and withdrawal. **(c)** Example of Monkey UL’s finger insertion into the left hole (left) and right hole (right).

**Supplementary figure 2. Task schemes and behavior results for each monkey.**

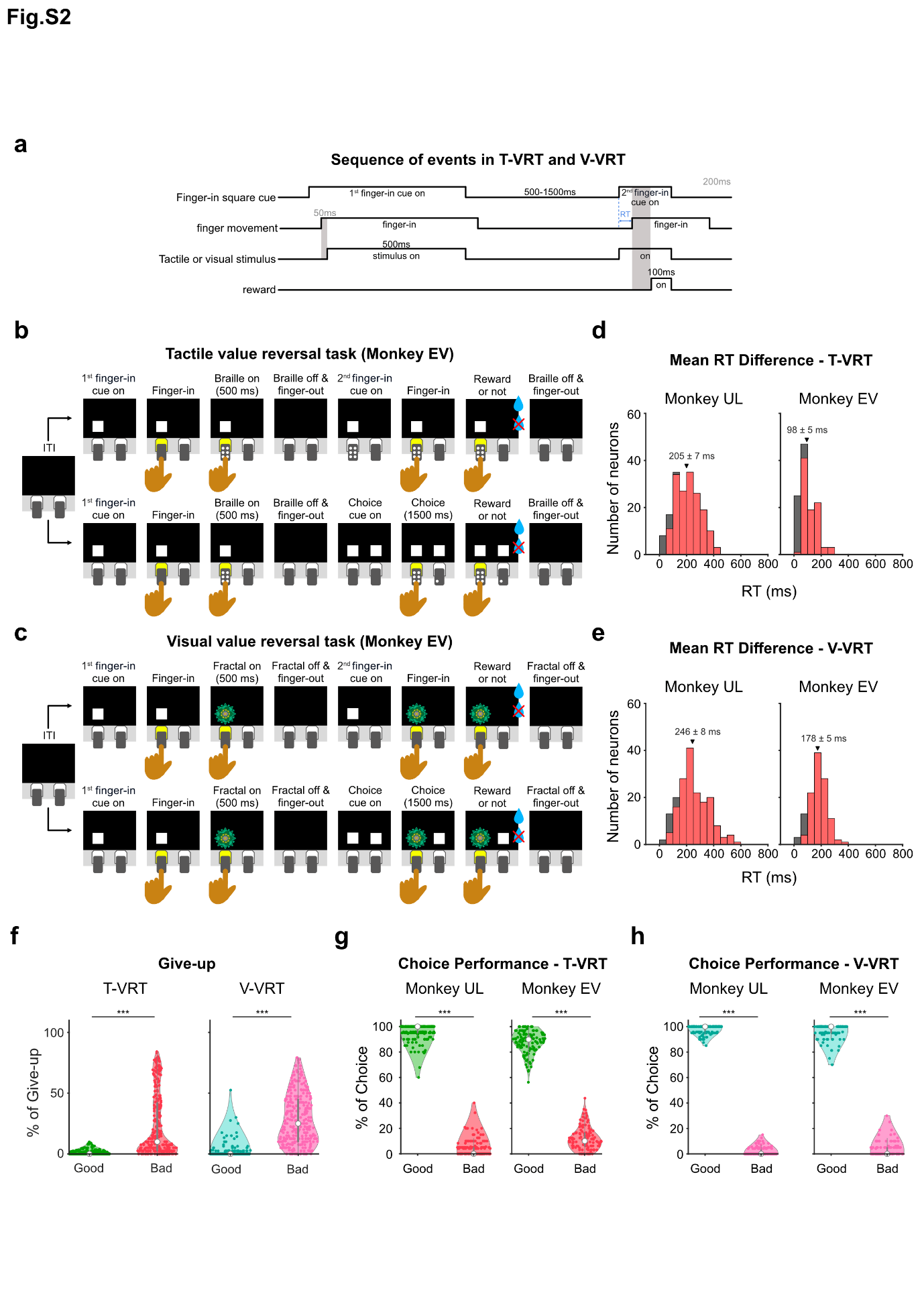
 **(a)** Sequence of events in a single trial of the T-VRT and V-VRT. **(b)** T-VRT for monkey EV. Instead of the colored cues used for monkey UL described in Figure 2c, only white square cues were used for monkey EV. In the choice trials for monkey EV, a single stimulus was presented before two white squares for choices appeared. **(c)** V-VRT for monkey EV. The V-VRT for monkey EV follows the same task procedure as the T-VRT, presenting fractal images instead of braille patterns. **(d)** Histogram of averaged differences in finger insertion reaction times for each monkey in the T-VRT (n = 180 for monkey UL and n = 119 for monkey EV). Red bars indicate neurons with statistically significant differences (two-tailed unpaired t-test, p < 0.05). Triangles and numerical values indicate the mean of the differences from all sessions (mean ± SE). **(e)** Histogram showing averaged differences in finger insertion reaction times for each monkey in the V-VRT (n = 180 for monkey UL and n = 119 for monkey EV). The format is consistent with (d). **(f)** Percentage of trials in which finger insertion was abandoned. Graphs indicate the percentage of trials in which the monkeys did not insert their fingers into the hole in response to the second finger-in cue presentation in the T-VRT (n = 299) and V-VRT (n = 299) (give-up trials). Paired t-testing was used to compare the percentages (***p < 0.005). **(g)** Performance in choice trials of the T-VRT. The percentages for selecting good and bad stimuli are plotted (n = 180 and n = 119 for monkeys UL and EV, respectively). The percentages of good and bad choices were compared by paired t-testing (***p = 1.000 x 10^-158^ for monkey UL, ***p = 8.851 x 10^-77^ for monkey EV). **(h)** Performance in choice trials of the V-VRT (n = 180 and n = 119 for monkeys UL and EV, respectively). The format is the same as in (g) (paired t-test, ***p = 8.069 x 10^-219^ for monkey UL, ***p = 1.897 x 10^-110^ for monkey EV).

**Supplementary figure 3. Location of value neurons in the putamen.**

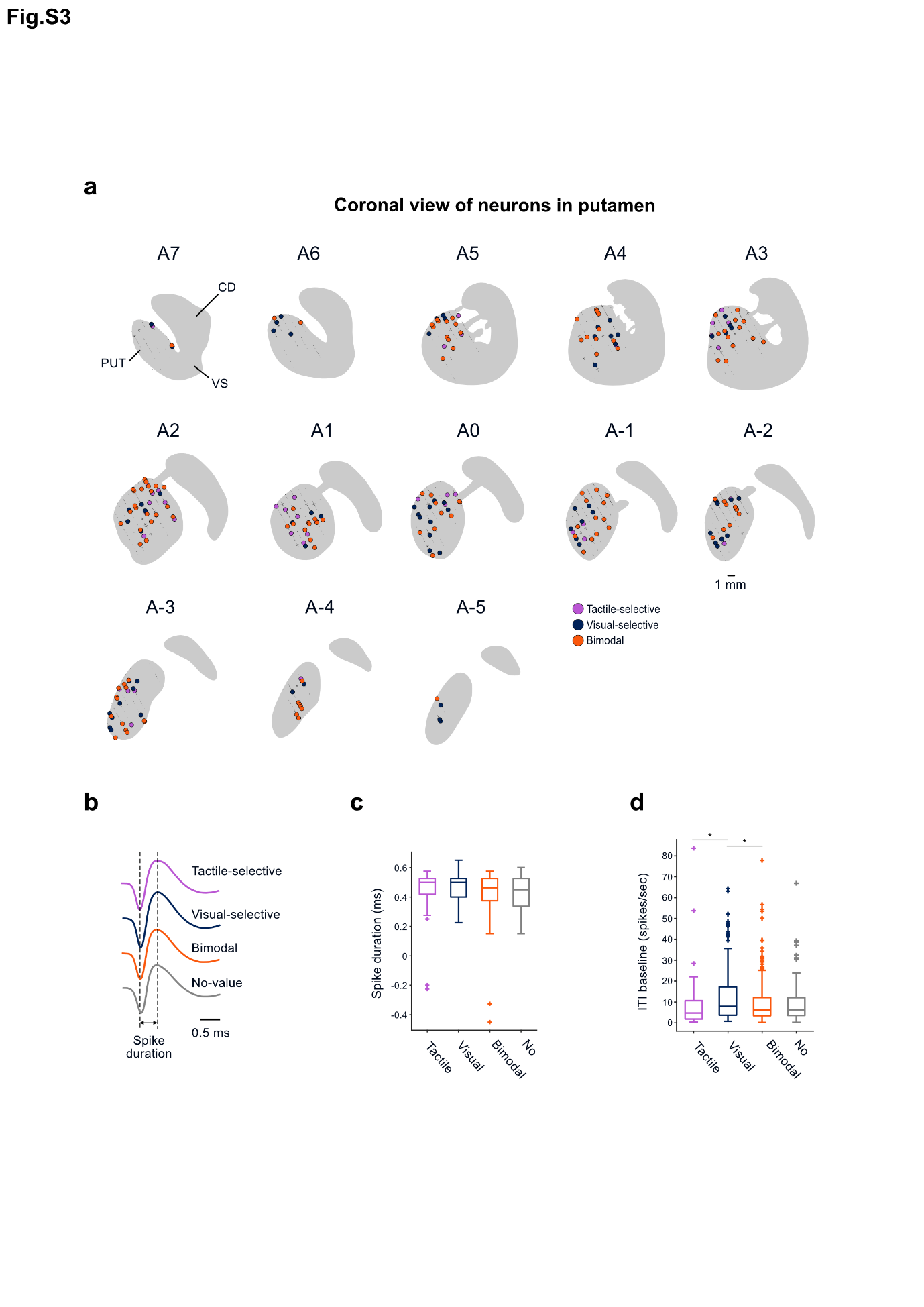
 **(a)** MR-based reconstruction of the recording sites of value neurons in coronal views of the putamen. AC numbers indicate distances from the anterior commissure (AC). Small dot: encountered neurons (*n* = 2257) during all the recording sessions. Cross: non-value neurons (*n* = 52). Purple circle: tactile-selective value neurons (*n* = 41). Navy circle: visual-selective value neurons (*n* = 77). Orange circle: bimodal value neurons (n = 129). **(b–d)** Electrophysiological properties of neurons recorded in the putamen. **(b)** Value-coding and non-value-coding neurons in the putamen had similar spike shapes. **(c)** Box plots comparing the spike durations between value-coding and non-value-coding neurons. The middle line indicates the median, and the bottom and top of the box correspond to the 25th and 75th percentiles, respectively. The upper and lower whiskers extend to the highest and lowest values within 1.5 times the interquartile range of the hinge. Outlying data beyond this are plotted as cross signs. A one-way ANOVA was applied to test the statistical significance. **(d)** Box plots comparing the baseline activities of value-coding and non-value-coding neurons during the inter-trial-interval. The baseline activities were compared by one-way ANOVA and post-hoc Bonferroni pairwise comparison (*p < 0.05).

**Supplementary figure 4. Analysis of differences in finger movements inside the hole.**

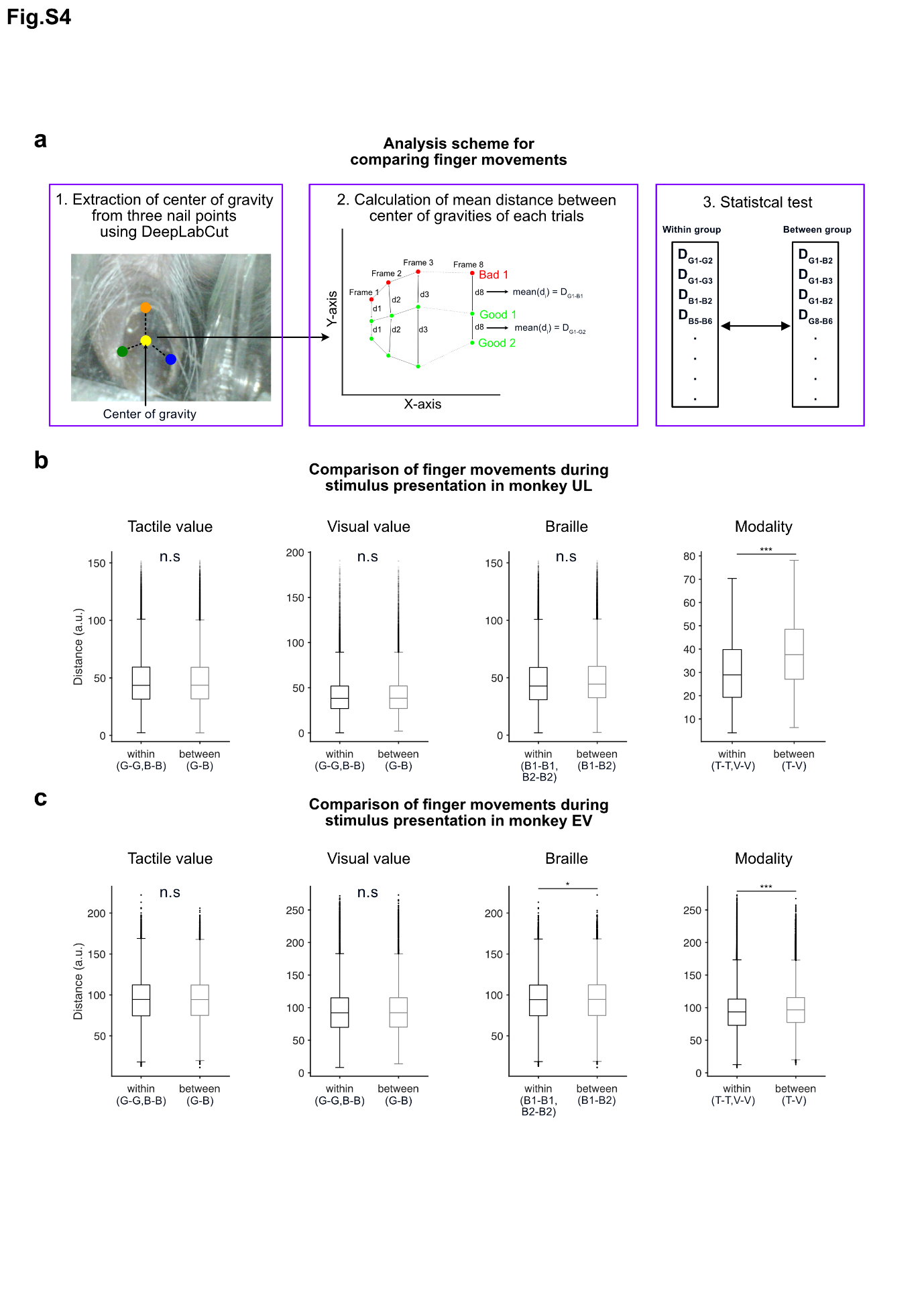
**(a)** Scheme for analyzing finger movements inside the hole. We determined the center of gravity for each tested frame using 3 nail points, extracted through DeepLabCut^43,44^ (left panel). Subsequently, traces of the center of gravity were plotted to calculate the distances among all combinations of the traces (middle panel). These distances were averaged, and the results were categorized based on their trial types. The statistical significance of differences in the finger trajectory distances was analyzed by two-tailed unpaired t-testing (p < 0.05) (right panel). **(b)** Box plots comparing the mean distances of the finger trajectories for monkey UL. The bottom and top of the box correspond to the 25th and 75th percentiles, respectively. The upper and lower whiskers extend to the highest and lowest values within 1.5 times the interquartile range of the hinge. The solid line inside each box plot represents the median of each group, and small dots outside the box plot indicate outliers. G: good stimuli. B: bad stimuli. B1: braille pattern 1. B2: braille pattern 2. T: T-VRT. V: V-VRT. Statistical significance was analyzed using two-tailed unpaired t-testing (p = 0.544 for tactile value, p = 0.252 for visual value, p = 0.538 for Braille pattern, ***p = 3.341 x 10^-52^ for modality). **(c)** Box plots comparing the mean distances of the finger trajectories for monkey EV. The format is the same as in (b). Statistically significant differences were observed in braille patterns and modality conditions (two-tailed unpaired t-testing, p = 0.472 for tactile value, p = 0.089 for visual value, *p = 5.510 x 10^-7^ for braille pattern, ***p=3.309 x 10^-77^ for modality).

**Supplementary figure 5. Positive and negative neuronal coding of tactile and visual value in the putamen.**

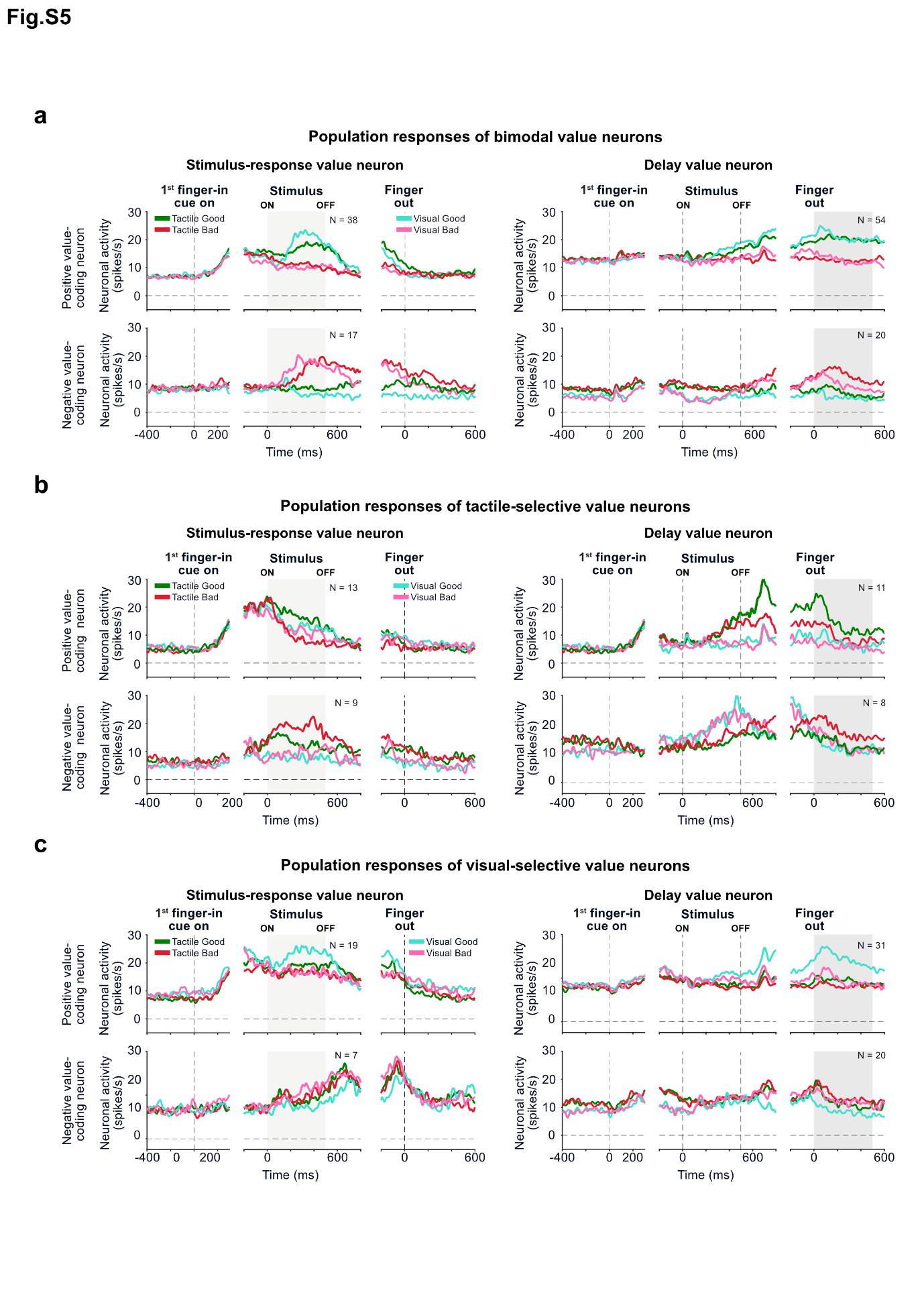
**(a-c)** Averaged neuronal responses are shown separately for neurons encoding positive values (positive value–coding neurons) and negative values (negative value–coding neurons). **(a)** Average responses to good and bad stimuli in bimodal value neurons. Green line: good tactile stimuli. Red line: bad tactile stimuli. Cyan line: good visual stimuli. Pink line: bad visual stimuli. **(b)** Average responses to good and bad stimuli in tactile-selective value neurons. The graph format is the same as in (a). **(c)** Average responses to good and bad stimuli in visual-selective value neurons. The graph format is the same as in (a).

**Supplementary figure 6. Changes in neural responses correlated with anticipated value-guided finger insertion across trials.**

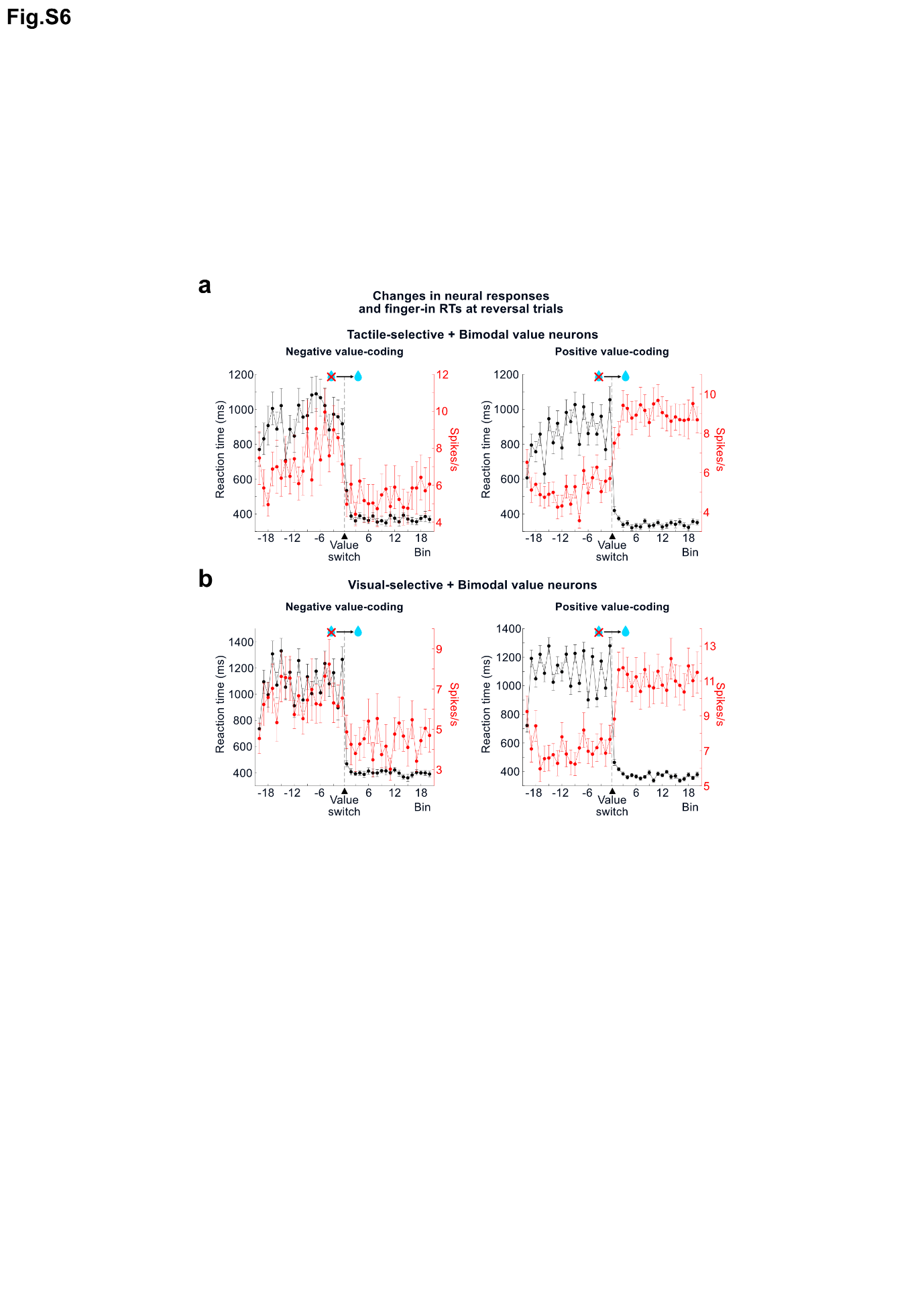

**(a)** Changes in neural responses and finger insertion speeds before and after value reversal in the T-VRT. The spike rates of value neurons (red) and finger-in reaction times (black) (mean ± SE) were plotted before and after the value reversal from bad to good. Neural spikes of tactile-selective value and bimodal value neurons were analyzed for the T-VRT. **(b)** Changes in neural responses and finger insertion speeds before and after value reversal in the V-VRT. The format is the same as in (a). Neural spikes of visual-selective value and bimodal value neurons were analyzed for the V-VRT.

**Supplementary figure 7. Correlation coefficient histograms for three types of value neurons in the T-VRT and V-VRT**.

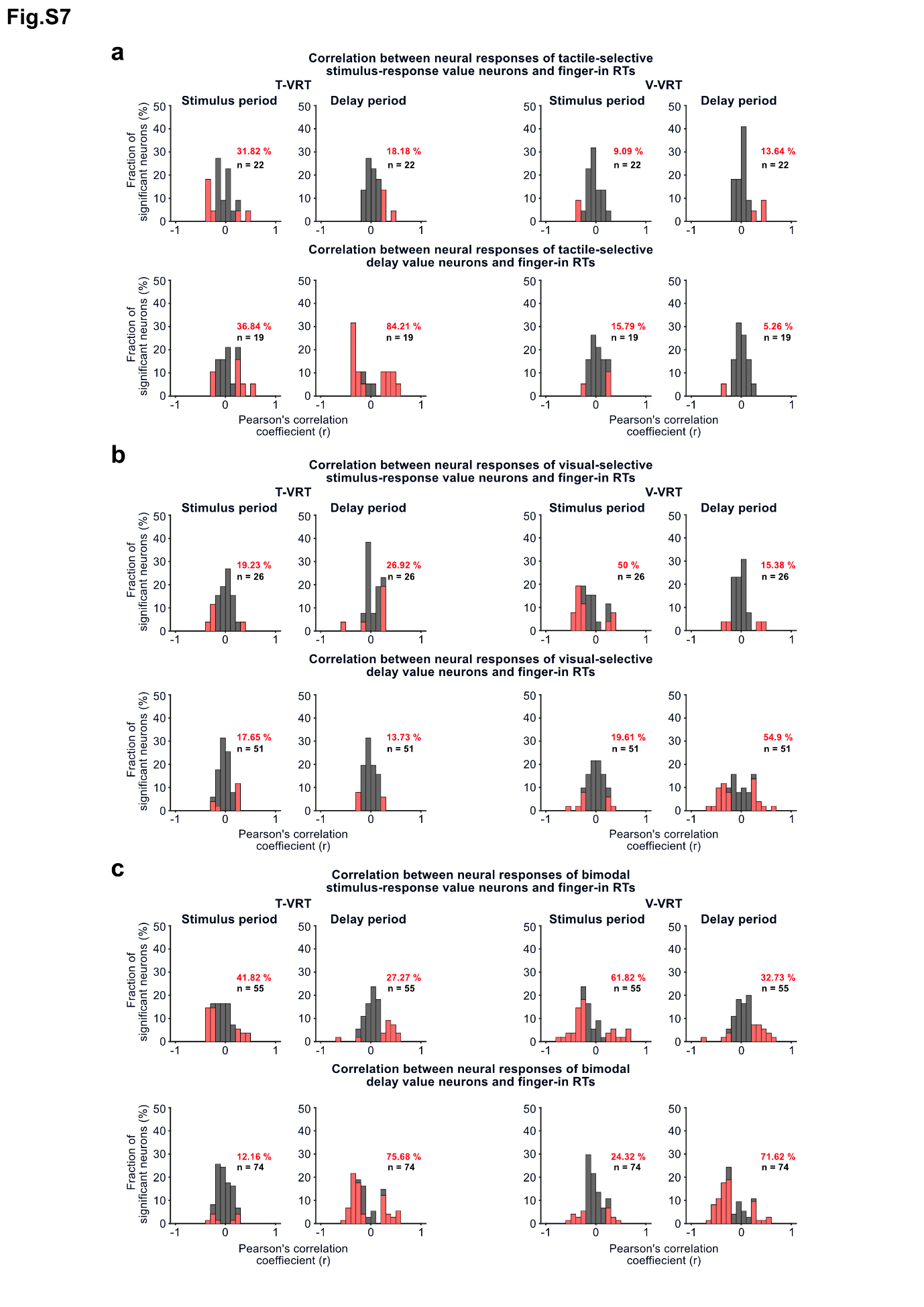

**(a)** Histograms of correlation coefficients for tactile-selective value neurons. Top panel shows the correlation coefficient histograms of tactile-selective value neurons responding during the stimulus presentation period in the T-VRT (left) and V-VRT (right). Bottom panel shows the histograms of tactile-selective value neurons responding during the blank delay period. Red bars indicate neurons with statistically significant correlations (Pearson correlation coefficient, p < 0.05). **(b)** Histograms of correlation coefficients for visual-selective value neurons. The format is the same as in (a). **(c)** Histograms of correlation coefficients for bimodal value neurons. The format is the same as in (a).

**Supplementary figure 8. Sensory modality decoding in parallel and convergent value processes using modality-selective and bimodal value neurons.**

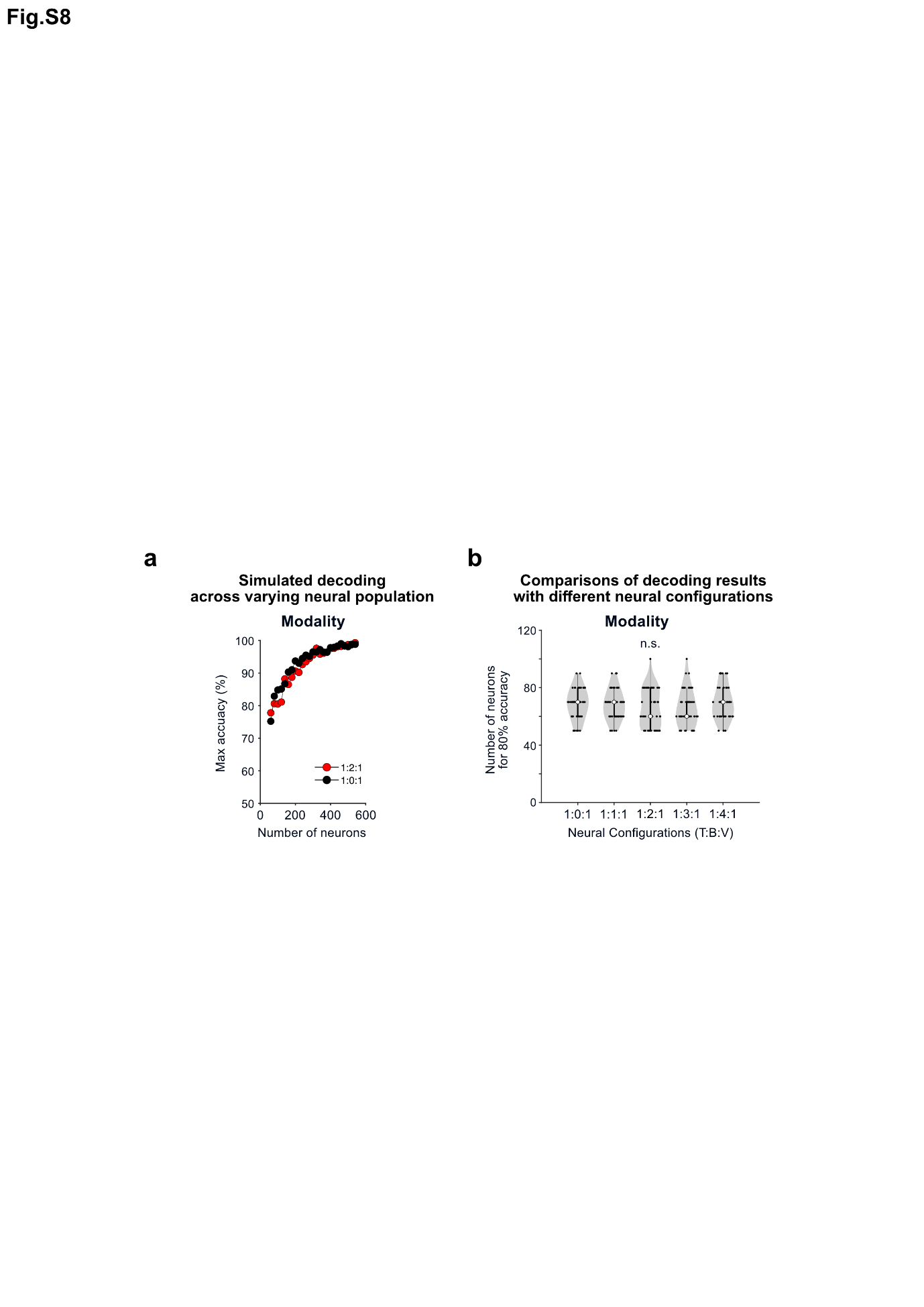

**(a)** Example plot showing the maximum values of modality decoding accuracy calculated with various total numbers of value neurons. Red and black dotted lines indicate convergent (1:2:1 ratio of tactile, bimodal, visual value neurons) and parallel (1:0:1) information processes, respectively. **(b)** Comparison of the minimum number of neurons required to achieve 80% modality decoding accuracy with 5 neural configurations representing parallel and convergent processes. A one-way ANOVA was applied to test statistical significance (F(4,245) = 1.28, p = 0.278).
